# Coexistent *PTEN* and *PIK3CA* alterations hyperactivate mTORC1 signaling in endometrial cancers and cause their selective sensitivity to mTORC1 inhibition

**DOI:** 10.64898/2026.02.12.705558

**Authors:** Hilla Solomon, Radha Mukherjee, Yu Chi Yang, Julia Meredith, Alison M. Schram, Sang Ah Yi, Xiaoping Chen, Michael Tribuzio, Harika Gundlapalli, Justin Meyerowitz, Elisa de Stanchina, Britta Weigelt, Heeseon An, Simon T. Barry, Jacqueline A. M. Smith, Mallika Singh, Neal Rosen

## Abstract

In approximately half of endometrial carcinoma (EC), PTEN loss-of-function and activating PI3K mutants coexist. Unlike cells with either single mutation, *PTEN*/*PIK3CA* coexistent alterations result in elevated membrane phosphatidylinositol (3,4,5)-trisphosphate (PIP3) levels and mTORC1 hyperactivation, rendering PI3K or AKT inhibition ineffective in blocking mTORC1 activity and tumor growth. The bi-steric mTORC1 kinase inhibitor, RMC-6272, suppresses mTORC1 activity and cell growth by reducing protein translation and cell cycle progression. *In vivo*, RMC-6272, but not PI3K inhibitors, effectively suppressed mTORC1 and growth of EC PDXs with coexistent *PTEN/PIK3CA* lesions. These findings are consistent with a phase I trial of bi-steric mTORC1 inhibitor RMC-5552, showing anti-tumor activity in patients with EC. PDXs with KRAS co-mutations regrew after RMC-6272 treatment, which was prevented by the addition of the RAS(ON) multi-selective inhibitor RMC-7977. Overall, these data suggest that mTORC1 hyperactivation drives ECs with coexistent *PTEN/PIK3CA* mutations, explain the limited antitumor activity of PI3K and AKT inhibitors, and support clinical evaluation of mTORC1 inhibitors as potential therapy for EC.

**Significance:** We have found the mechanistic consequences of *PTEN/PIK3CA* co-alterations in endometrial tumors and that these mutations result in a profound hyperactivation of mTORC1 signaling. Single mutant tumors are sensitive to PI3K inhibition but those with both mutations are insensitive to PI3K or AKT inhibition but are exquisitely dependent on mTORC1 kinase. This provides strong preclinical rationale for targeting mTORC1, alone or combined with RAS inhibition (in RAS co-mutant tumors), as an effective therapeutic strategy.

## Introduction

Endometrioid uterine carcinomas have a uniquely high frequency of coexistent *PTEN*-inactivating lesions and activating mutations of *PIK3CA* ^1^, leading to high levels of PI3K pathway output. The *PIK3CA* gene encodes the p110α catalytic subunit of the phosphatidylinositol 3-kinase (PI3K) Class I lipid kinase which catalyzes the formation of phosphatidylinositol-3,4,5-triphosphate (PIP3). The catalytic subunit binds to the p85 regulatory subunit (encoded by *PIK3R1* gene) which reduces its catalytic activity. Activated receptor tyrosine kinases (RTKs) relieve p85 inhibition of the p110α catalytic subunit and enhance synthesis of PIP3 ^2^. PIP3 binds to the pleckstrin homology (PH) domain of a set of proteins including AKT and PDK1 causing them to bind to membranes and activate substrates that regulate proliferation, cell survival, metabolism, and other processes. The mammalian target of rapamycin (mTOR) is downstream of PI3K/AKT and is a major regulator of cell growth integrating energy, nutrient and growth factor supply by controlling cap-dependent translations ^3^. Termination of the PIP3 signal is mediated by lipid phosphatases that convert PIP3 into PIP2. This includes the Phosphatase and Tensin homolog (PTEN) that removes a phosphate group from the 3-position of PI(3,4,5)P3, converting it to PI(4,5)P2. This action terminates the PIP3 signal and acts as a negative regulator of the PI3K pathway ^4^.

Homeostatic regulation of the PI3K/AKT/mTOR pathway is maintained by a complex network of negative feedback mechanisms. Feedback inhibition of upstream signaling is induced by activated AKT or mTORC1, which inhibit the expression or activation of some RTKs. This upstream feedback is overcome in part by constitutive activation of signaling by PI3K or AKT activating mutants ^5^. PI3K signaling is also regulated downstream by PI3K/mTOR dependent induction of PTEN translation and expression ^6^. mTORC1 regulates the cap-dependent translation of PTEN, thus activation of PI3K/AKT/mTOR signaling increases PTEN protein expression and blunts the amplitude and duration of the PI3K signal via negative feedback. By contrast, inhibition of PI3K signaling or mTORC1 activity, reduces PTEN expression disrupting the negative feedback and buffering the reduction in pathway output ^6^.

Dysregulation of PI3K signaling is a common event in solid tumors and is most often due to gain of function mutations of the catalytic or regulatory subunits of class 1 PI3K, amplification of the gene encoding HER2, or inactivating mutations of PTEN. Endometrial carcinomas (EC) have a higher frequency of PI3K/mTOR pathway alterations than any other cancers (*PTEN* 67%, *PIK3CA* catalytic subunit 57%, *PIK3R1* regulatory subunit 34% (Supplementary Fig. 1A) ^7–9^. Moreover, *PTEN* inactivating mutations coexist with *PIK3CA* catalytic mutations in 30% of these tumors, a far greater prevalence than any other tumor (Supplementary Fig. 1B). The reason for this is not understood but the data suggest that activation of PI3K signaling plays an important role in EC tumor development. Nevertheless, PI3K/AKT/mTOR pathway inhibitors are not used in the treatment of patients with EC ^10–12^. The effects of rapamycin analogs on these tumors have been tested but only a modest clinical benefit was observed ^11,13–16^.

We hypothesized that the coexistence of *PTEN* loss and activating *PIK3CA* mutants would disable both upstream (reduction in PI3K signaling) and downstream (increase in PTEN expression) feedback mechanisms and result in very high mTORC1 output that drives tumor growth and is insufficiently inhibited by agents such as rapalogs, PI3K or AKT inhibitors. In this report, we show that EC models with both mutants have elevated membrane-bound PIP3 and phosphorylation of mTORC1 kinase substrates. Neither PI3K nor AKT inhibitors effectively suppressed mTORC1 kinase activity, nor did they arrest tumor growth *in vivo*. In contrast, because these double mutant tumor cells are exquisitely dependent on mTORC1 activity, the selective mTORC1 kinase inhibitor, RMC-6272, effectively blocked the phosphorylation of mTORC1 targets and inhibited the growth of these tumors.

## Results

### The sensitivity of EC tumor cells to PI3K or AKT inhibition is reduced in cells with coexistent *PTEN* inactivating and *PIK3CA* activating mutants

Previously we had observed that the *PIK3CA/PTEN* double mutant EC cell lines have relatively higher mTORC1 activity as shown by increased phosphorylation of 4E-BP1 and S6K as compared to single *PIK3CA* or *PTEN* mutant cell lines (6). We sought to assess the response of a panel of six EC cell lines, each with a distinct PTEN and PI3K mutation status, to selective inhibitors targeting different nodes of the pathway, with the aim of delineating their signaling and growth vulnerabilities. These include KLE (WT *PTEN* /WT *PIK3CA*), MFE280 (WT *PTEN/*mutant *PIK3CA^H1047Y,I391M^*), AN3CA (mutant *PTEN^R130fs^ /*WT *PIK3CA*), and three cell lines in which both *PIK3CA* and *PTEN* are mutated: MFE296 (mutant *PTEN^R130Q,N323fs^ /* mutant *PIK3CA^P539R,I20M^*), HEC6 (mutant *PTEN^V290fs,V85fs^ /* mutant *PIK3CA^R108H,C420fs^*) ^17^, and high-grade uterine carcinoma PDX-derived cells (mutant *PTEN^M134del,R173H^/* mutant *PIK3CA^E542K^*). AN3CA and HEC6 cell lines have PTEN frameshift mutations and undetectable PTEN protein expression. MFE296 and the EC PDX-derived cells carry PTEN missense mutations, which impair their phosphatase activity ^18,19^. The high-grade uterine carcinoma PDX-derived cells carry the *PIK3CA* helical domain hot spot mutation (E542K). MFE280, MFE296 and HEC6 carry two *PIK3CA* mutations in cis, which increase PI3K specific activity and oncogenicity ^20^ (Supplementary Table 1).

To fully inhibit PI3K, we combined the selective PI3K-α and PI3K-β inhibitors (BYL719 and AZD8186, respectively) at concentrations at which they are selective for inhibition of PI3K ^21^. PI3K inhibition significantly blocked PI3K/AKT signaling in all cell lines, as shown by a rapid reduction in phosphorylated AKT and its targets GSK3β and PRAS40. However, mTORC1 activity, assessed by the phosphorylation of 4E-BP1, ULK1, S6K, and S6, was more effectively inhibited in cells expressing wild-type *PTEN/PIK3CA* (KLE) or single mutations (AN3CA or MFE280) than in those with coexistent *PTEN* and *PIK3CA* alterations (MFE296, HEC6, high-grade uterine carcinoma PDX-derived cells) (Fig. 1A-D, Supplementary Fig. 2, upper panels). For example, the phosphorylation of 4E-BP1(S65) declined 1h after treatment, reaching 1.5% of initial levels by 24 hours in MFE280 cells, 35.5% in AN3CA cells and 13.5% in KLE cells. However, in the double mutant cells phosphorylation of 4E-BP1 (S65) was less affected by PI3K inhibition, reaching 57% of initial levels by 24 hours in MFE296, or 89% in HEC6 or the PDX-derived cells. This suggests that with PI3K inhibition, mTORC1 activity was inhibited but remained higher in *PTEN/PIK3CA* mutant coexisting cells as compared with cells bearing only single mutations. The proliferation of tumor cells with coexistent *PTEN* and *PIK3CA* alterations was relatively insensitive to PI3K inhibition, while the proliferation of cells with wild-type *PTEN/PIK3CA* or either *PTEN* or *PIK3CA* single mutations was significantly inhibited by PI3K inhibitors (Fig. 1A-D and supplementary Fig. 2, lower panels).

**Figure 1:**
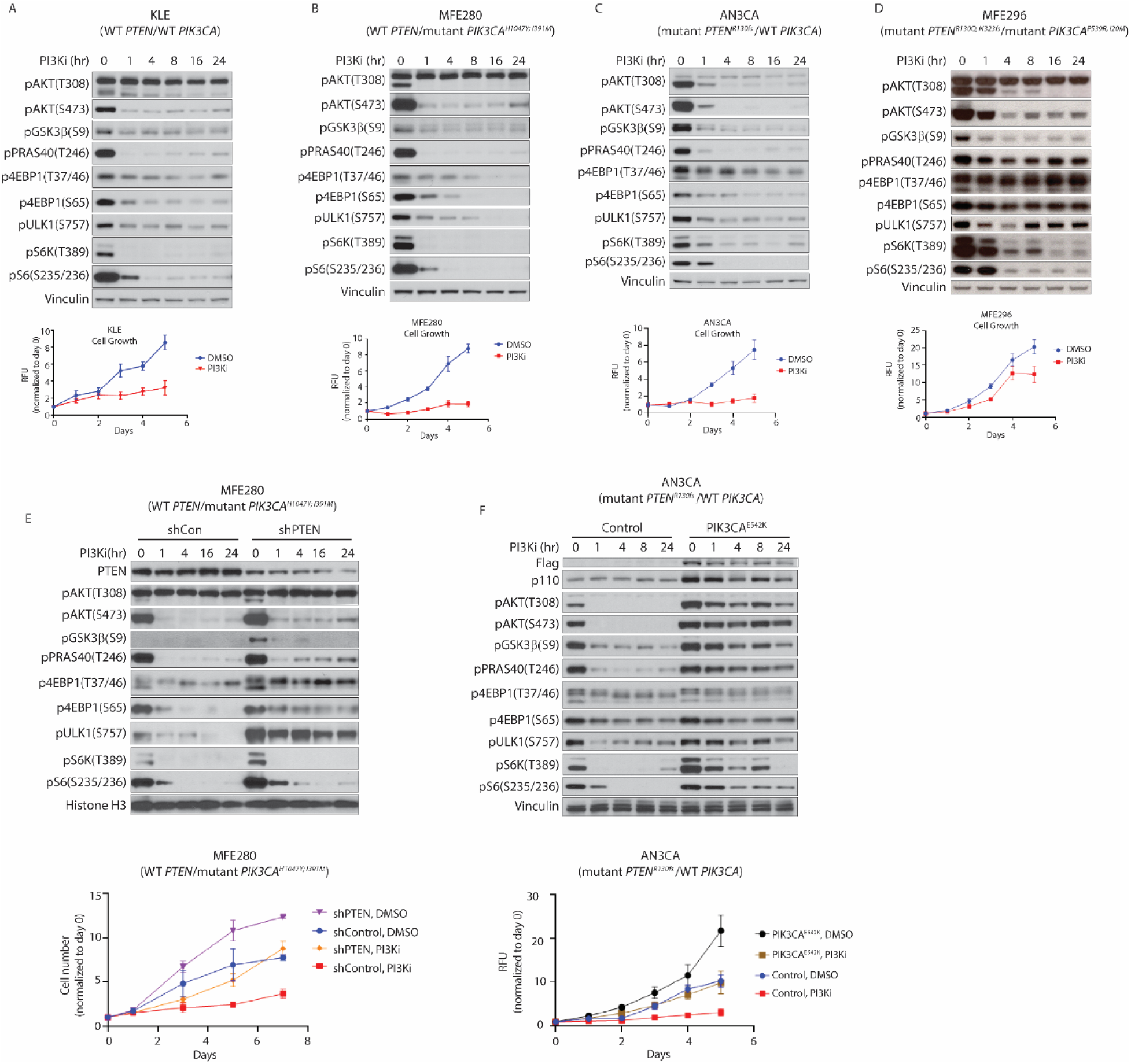
The sensitivity of EC cells to PI3K inhibition is reduced in cells with coexistent *PTEN* inactivating and *PIK3CA* activating mutants. The following EC cell line models were treated with combination of 1μM BYL-719 + 250 nM AZD8186 (PI3Ki) for the indicated time points: (A) KLE (WT *PTEN/* WT *PIK3CA*), (B) MFE280 (WT *PTEN/*mutant *PIK3CA^H1047Y; I391M^*), (C) AN3CA (mutant *PTEN^R130fs^ /*WT *PIK3CA*) and (D) MFE296 (mutant *PTEN^R130Q, N323fs^ /*mutant *PIK3CA^P539R, I20M^*). (E) MFE280 cells were stably infected with shRNA against PTEN. (F) AN3CA cells were transiently transfected with the mutant *PIK3CA^E542K^* for 24 hours prior to treatment. Upper panels: Immunoblots depicting PI3K/mTOR signaling output. Lower panels: cell proliferation measured by alamarBlue assay (A-D, F) or by direct cell counting using trypan blue staining and hemocytometer (E).

Overall, these results show that mTORC1 activity is associated with the proliferation of EC cells with coexistent *PTEN* and *PIK3CA* alterations, and that PI3K inhibition is insufficient to completely block mTORC1 activity and the proliferation of these double mutant EC cells.

We further tested this hypothesis in isogenic EC models in which we controlled *PTEN* or *PIK3CA* expression. MFE280 cell line endogenously expresses WT *PTEN* and mutant *PIK3CA^H1047Y;I391M^*. PTEN knockdown increased phosphorylation of AKT, its target GSK3β, and mTORC1 substrates ULK1, S6K, S6 and 4E-BP1(T37,46) (Fig. 1E, upper panel). In addition, we found that the basal rate of proliferation increased in the double mutant cells as compared to the cells with mutant *PIK3CA* alone (Fig. 1E, lower panel). PI3K inhibition reduced phosphorylation of AKT and mTORC1 substrates and inhibited proliferation in controls with WT *PTEN* expression. In contrast, PI3K activity in the double mutant cells was inhibited but showed a greater rebound (pAKT, pRAS40, pGSK3b) and there was no inhibition of mTORC1 substrates p4E-BP1 (T37,46) and pULK (Fig. 1E, upper panel). The double mutant cells proliferated faster than the WT *PTEN* cells in the context of PI3K inhibition (Fig. 1E, lower panel).

A second isogenic system was developed from AN3CA cells with WT *PIK3CA* and the *PTEN^R130Qfs^* frame-shift mutation, where the helical domain mutant *PIK3CA^E542K^* was expressed, creating an isogenic system with a coexistent PI3K activating mutant and no PTEN function (Fig. 1F, upper panel). Introduction of the *PIK3CA^E542K^*mutant increased phosphorylation of AKT and the mTORC1 substrate S6K. Changes in 4E-BP1 phosphorylation were marginal in the double mutant compared to the control cells, suggesting sufficiency of PTEN loss of function alone for increase in 4E-BP1 phosphorylation in this model. Strikingly, upon treatment with PI3K inhibitors, the reduction of pAKT, its substrates (pGSK3β, pPRAS40), pS6K, and pS6 was markedly less pronounced in the double mutant cells as compared to that in control cells with PTEN mutation alone, while p4EBP1 remained comparably sensitive to PI3K inhibitors in both the single and double mutant cells. Like the MFE280 isogenic system, the basal proliferation of the AN3CA cells increased with the expression of *PIK3CA* mutation and these cells continued to grow more rapidly in the presence of PI3K inhibition as compared to the single mutant control cells (Fig. 1F, lower panel).

Taken together, our findings suggest that *PTEN* loss-of-function in the context of *PIK3CA* mutation enhances mTORC1 activity as evidenced by the marked increase in the phosphorylation of 4EBP1 and ULK1, consistent with PTEN being an upstream suppressor of mTORC1 ^6^. Phosphorylation of 4EBP1 and ULK1 become resistant to PI3K inhibition in these double mutant cells (Fig. 1E). On the other hand, subsequent overexpression of gain-of-function mutant *PIK3CA* in cells with *PTEN* loss leads to the activation of pAKT and pS6K (Fig. 1F). Nonetheless, coexistent *PTEN* loss and *PIK3CA* mutation results in sustained mTORC1 activity in both isogenic models, which blunts the ability of PI3K inhibitors to block PI3K/AKT/mTORC1 signaling, leading to a relatively reduced impact on growth inhibition.

Since PI3K activates downstream signaling by inducing PIP3 formation, and PTEN opposes this by dephosphorylating PIP3, we asked whether cells with the double mutations have increased PIP3 levels. PIP3 levels were measured using an intracellular reporter assay ^22^. In this assay, GFP is fused to the PH domain of AKT (GFP-PH^AKT^) which binds PIP3 in the membrane. Quantifying translocation of GFP-PH^AKT^ to the plasma membrane allows assessment of PIP3 levels. Our data revealed that membrane PIP3 was significantly elevated in the MFE296 (mutant *PTEN^R130Q, N323fs^/*mutant *PIK3CA^P539R, I20M^*) cell line with a mean value of 1.094, compared with MFE280 (WT *PTEN/*mutant *PIK3CA^H1047Y, I391M^*) with a mean value of 0.6935 (p-value < 0.001) and AN3CA (mutant *PTEN^R130fs^/*WT *PIK3CA*) with a mean of 0.9306 (p-value < 0.001) (Supplementary Fig. 3A and C). This suggests that while *PTEN* loss alone increases membrane associated PIP3 levels as compared to single mutant *PIK3CA* cells, dual alterations of *PIK3CA* and *PTEN* increase membrane associated PIP3 the most. We compared PIP3 in control AN3CA cells (mutant *PTEN^R130fs^* /WT *PIK3CA/*) with those transfected with mutant *PIK3CA^E542K^* and found an increase in the mean value from 0.9233 in the control cells to 1.172 in the double mutants (mutant *PTEN^R130fs^ /*mutant *PIK3CA^E542K^*) (p-value <0.001) (Supplementary Fig. 3B-C), similar to PIP3 levels observed above in the double mutant MFE296 cell line. Across both cell lines and the isogenic model, *PTEN* lesions together with *PIK3CA* mutations lead to elevated membranous PIP3 and increased activity of downstream AKT and mTORC1 signaling, which likely contributes to reduced sensitivity to PI3K inhibitors.

Phosphoinositide-dependent protein kinase 1 (PDK1) binds to PIP3 in the membrane via its PH domain, leading to its autophosphorylation on serine 241 and its activation ^23^. Basal pPDK1 levels were 2-fold higher in the *PTEN/PIK3CA* double mutant MFE296 cells compared with single *PTEN* or *PIK3CA* mutant cells. After PI3K inhibition, PDK1 phosphorylation rebounded to much higher levels in the MFE296 double mutant cells, reaching 75% of initial levels at 24 hours as compared to cells with *PTEN* loss alone (AN3CA) or *PIK3CA* mutations alone (MFE280), which reached 28% or 38% of initial levels at 24-hours, respectively (Supplementary Fig. 4A-D). This is consistent with higher levels of PIP3 in the membrane in double mutant cells. In both isogenic models, phosphorylated PDK1 levels were higher at 24 hours of PI3K inhibition in the *PTEN/PIK3CA* coexistent mutation cells, as compared with the single mutants (Supplementary Fig. 4E-F). Since PDK1 phosphorylation at S241 depends on PIP3 binding ^23^, these findings support the conclusion that PIP3 levels are elevated in *PTEN/PIK3CA* double mutant cells, which consequently contribute to hyperactivation of the PI3K/AKT/mTOR pathway.

PI3K exerts its oncogenic functions by regulating various downstream effectors, including the AKT/mTOR signaling pathway ^24,25^. We asked whether the resistance of *PTEN/PIK3CA* double mutant cells to PI3K inhibitors is due to hyperactivated AKT. The *PTEN/PIK3CA* double mutant cell line, MFE296 (mutant *PTEN^R130Q, N323fs^*/mutant *PIK3CA^P539R, I20M^*), was treated with either the allosteric AKT1/2/3 inhibitor, MK2206, or with the pan-AKT kinase ATP-competitive inhibitor, AZD5363 (capivasertib) (Fig. 2A-B). The effects of the AKT inhibitors and PI3K inhibitors on cell growth and PI3K/mTOR signaling output were very similar. The allosteric inhibitor, MK2206, binds near the N-terminal AKT and prevents its binding to the membrane and its phosphorylation and activation by PDK1 and mTOR ^26^. AZD5363 is an ATP-competitive inhibitor of AKT kinases that induces feedback reactivation of RTK/PI3K signaling and thus induces AKT(T308) and AKT(S473) ^27^. Both types of inhibitors reduce phosphorylation of the AKT substrates GSK3β and PRAS40. However, neither inhibits the phosphorylation of mTORC1 targets 4E-BP1 and ULK1 very well, nor do they inhibit the *in vitro* proliferation of the double mutant MFE296 cells (Fig. 2A-B). Moreover, MK2206 inhibited the phosphorylation of 4E-BP1, ULK1, S6K and S6 in MF280 (WT *PTEN/*mutant *PIK3CA^H1047Y; I391M^*), but in the MFE280 cells, in which *PTEN* was knocked-down, the phosphorylation of these mTORC1 substrates was less sensitive to MK2206 (Fig. 2C). Like the PI3K inhibitors, MK2206 inhibited the proliferation of MFE280 control cells more effectively than that of double-mutant MFE280 isogenic cells (Fig. 2D). Similar data were obtained in the AN3CA isogenic model (Fig. 2E-F, Supplementary Fig. 5). Taken together, our findings suggest that EC cells harboring concurrent *PTEN* and *PIK3CA* alterations display elevated PIP3 levels and enhanced overall mTORC1 kinase activity, making it difficult for PI3K or AKT inhibitors, which only target select aspects, to effectively block the pathway. We therefore hypothesize that a direct inhibition of mTOR will be more effective in inhibiting the pathway and cell growth.

**Figure 2:**
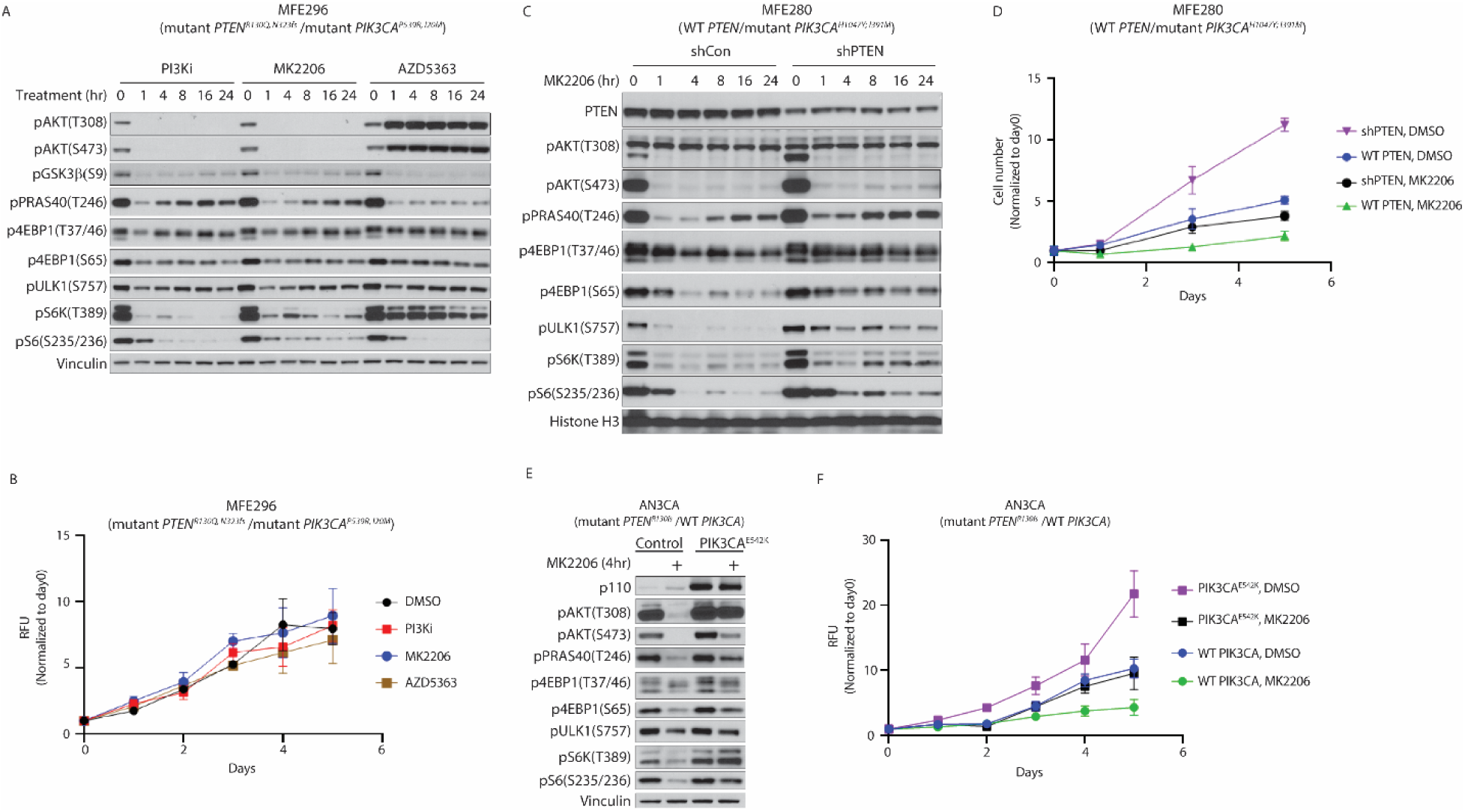
The sensitivity of EC cells to AKT inhibition is reduced in cells with coexistent *PTEN* inactivating and *PIK3CA* activating mutants. (A-B) MFE296 (mutant *PTEN^R130Q, N323fs^ /*mutant *PIK3CA^P539R, I20M^*) cells were treated with either 1μM BYL-719 + 250 nM AZD8186 (PI3Ki), 1μM MK2206 or 1μM AZD5363 for the indicated time points. (A) Immunoblots depicting PI3K/mTOR signaling output. (B) cell proliferation measured by alamarBlue assay. (C-D) MFE280 (WT *PTEN/*mutant *PIK3CA^H1047Y; I391M^*) cells were stably infected with shRNA against PTEN and treated with 1 μM MK2206. (C) Immunoblots depicting PI3K/mTOR signaling output. (D) Cell proliferation measured by direct cell counting using trypan blue staining and hemocytometer. (E-F) AN3CA (mutant *PTEN^R130fs^ /*WT *PIK3CA*) cells were transiently transfected with mutant *PIK3CA^E542K^*. (E) After 24 hours cells were treated with 1 μM MK2206 for 4 hours and immunoblots depicting PI3K/mTOR signaling output are shown. (F) After 24 hours of transfection with mutant *PIK3CA^E542K^*, cells were treated with 1 μM MK2206 for the indicated time points, and cell proliferation was assessed by alamarBlue assay.

### mTOR kinase inhibitors suppress mTOR activation and block the proliferation of EC cells with coexistent *PIK3CA* and *PTEN* alterations

We tested this idea with each of two mTOR kinase inhibitors, AZD8055, which inhibits mTORC1 and mTORC2, and a selective inhibitor of mTORC1 kinase, RMC-6272 ^28^. mTORC2 kinase phosphorylates AKT S473 and its inhibition is strongly associated with hyperglycemia, which limits the therapeutic potential of pan-mTOR kinase inhibitors, whereas RMC-6272 is an mTORC1 selective inhibitor and does not cause hyperglycemia in preclinical models. RMC-6272 is a preclinical tool compound representative of the clinical investigational agent RMC-5552, with the two compounds exhibiting comparable *in vitro* and *in vivo* properties ^28^. We also tested rapamycin, an allosteric inhibitor of mTOR which inhibits mTORC1 more potently than mTORC2 but with limited effect on levels of p4E-BP ^29^.

MFE296 proliferation was almost completely blocked by AZD8055 but almost not affected by PI3K inhibition or rapamycin (Fig. 3A). AZD8055 potently inhibited the phosphorylation of mTOR substrates including 4E-BP1, AKT(T308) and AKT(S473). Neither PI3K inhibition nor rapamycin inhibited 4E-BP1 phosphorylation, but rapamycin inhibited S6K phosphorylation (Fig. 3B). RMC-6272 potently inhibited the phosphorylation of mTORC1 substrates, including 4E-BP1, within 4 hours of treatment in MFE296 cells (Fig. 3D). As previously reported, the phosphorylation of mTORC2 target AKT(S473) was unaffected. pAKT(T308) was induced by AZD8055 and RMC-6272 due to relief of RTK/PI3K negative feedback ^22^.

**Figure 3:**
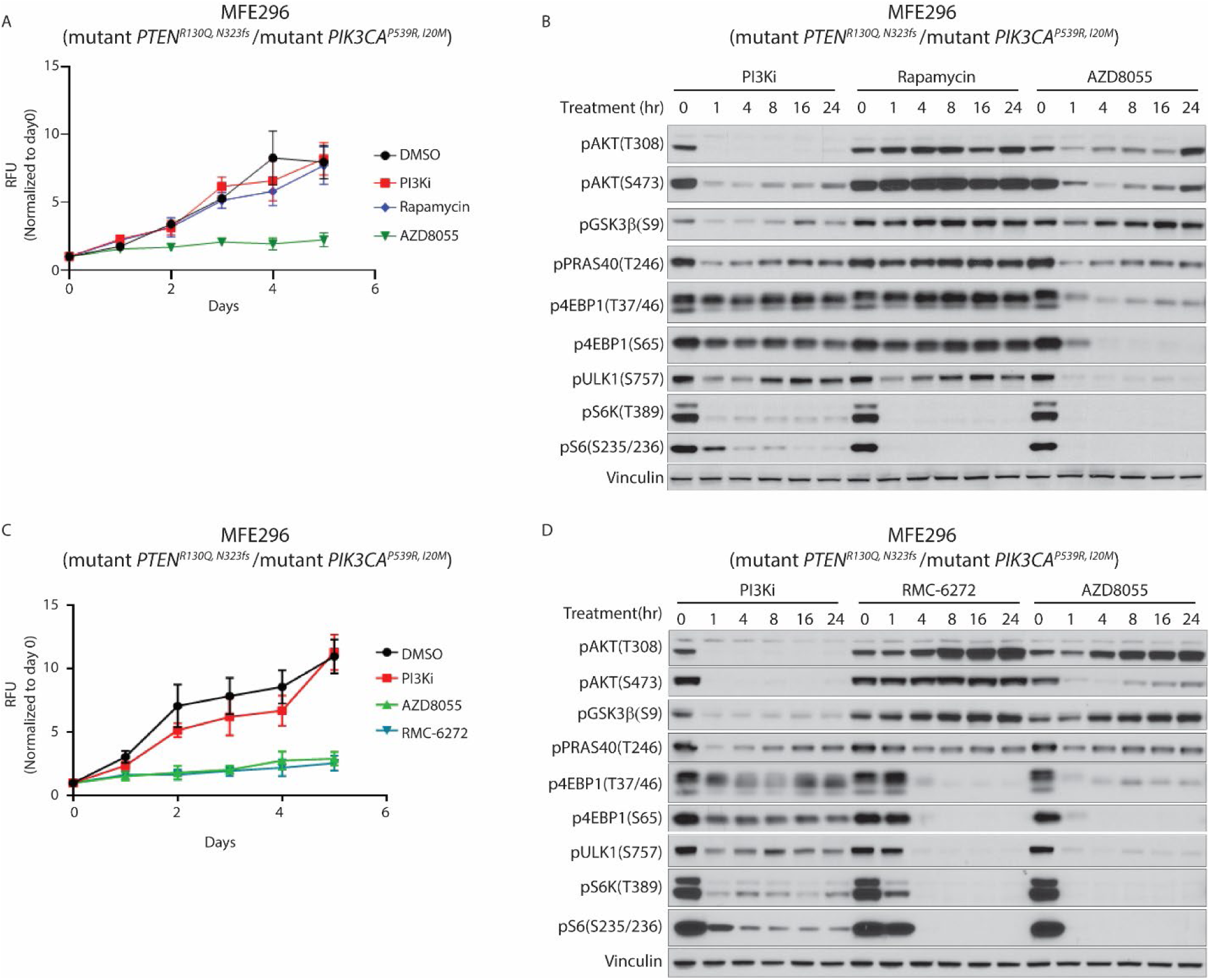
*PTEN/PIK3CA* double mutant EC cells are dependent on mTORC1. MFE296 (mutant *PTEN^R130Q, N323fs^ /*mutant *PIK3CA^P539R, I20M^*) cells were treated with the following inhibitors: a combination of 1μM BYL-719 and 250 nM AZD8186 (PI3Ki), 500 nM AZD8055(A-D), 50 nM rapamycin (A-B), or 500 pM RMC-6272 (C-D) for the indicated time points. (A and C) Cell growth was assessed by alamarBlue assay. (B and D) PI3K/mTOR signaling output was measured by immunoblotting using specific antibodies.

RMC-6272 and AZD8055 had comparable growth inhibition effects on MFE296 (*PTEN/PIK3CA* double mutant) EC cells, suggesting that inhibition of mTORC1 kinase is sufficient for effective growth inhibition (Fig. 3C). Similar results were obtained in another *PTEN/PIK3CA* double mutant cell line, HEC6 (mutant *PTEN^V290fs,V85fs^ /* mutant *PIK3CA^R108H,C420fs^*), and in high-grade uterine carcinoma PDX-derived cells (mutant *PTEN^M134del,R173H^ /* mutant *PIK3CA^E542K^*) (supplementary Fig. 6).

### mTORC1 inhibition reduces proteins required for G0/G1 transition

We then explored the mechanism whereby mTORC1 inhibitors block EC cell proliferation. mTORC1 activates mRNA translation by phosphorylating the eIF4E-binding protein, 4E-BP1 and ribosomal S6 kinases (S6K), thereby promoting the expression of proteins required for G1 progression, including cyclins, CDKs, and MYC ^30,31^. We measured the effect of PI3K or mTORC1 kinase inhibition on protein translation with the puromycin labeling assay. In this assay, puromycin, which mimics an aminoacyl-tRNA molecule, is incorporated into nascent peptide chains and the incorporation of puromycin is detected with a specific anti-puromycin antibody ^32^. Cells were treated with either PI3K inhibitors (BYL719 + AZD8186) or RMC-6272 for 24 hours, followed by a 30 minutes puromycin (1μM) pulse. In double mutant MFE296 cells, PI3K inhibition caused a modest reduction in puromycin incorporation as compared to DMSO control, whereas RMC-6272 treatment led to ∼5.5-fold decrease in 24 hours than DMSO control and significantly lower than PI3K inhibitors treatment (Fig. 4A). In contrast, in MFE280 (WT *PTEN/*mutant *PIK3CA^H1047Y; I391M^*) cells, both PI3K inhibitors or RMC-6272 caused a comparable reduction in protein synthesis at 24 hours (Fig. 4B). These data are consistent with the greater effect of PI3K inhibitors on 4E-BP1 phosphorylation in the single mutant MFE280 cells than in the double mutant MFE296 cells (Fig. 1B and D), and also consistent with mTORC1 inhibitor showing stronger inhibition of 4E-BP1 phosphorylation than the PI3K inhibitors in the double mutant cell line. Basal protein synthesis of AN3CA mutant *PTEN^R130fs^/*mutant *PIK3CA^E542K^* cells was slightly higher than in the WT *PIK3CA* control cells. Although the degree of inhibition of protein synthesis by PI3K inhibitors was similar in the double mutant and the single mutant AN3CA cells, overall protein synthesis levels was slightly higher in the former (Fig. 4C). Strikingly, RMC-6272 treatment led to a profound inhibition in protein synthesis in both the single and double mutant isogenic cells.

**Figure 4:**
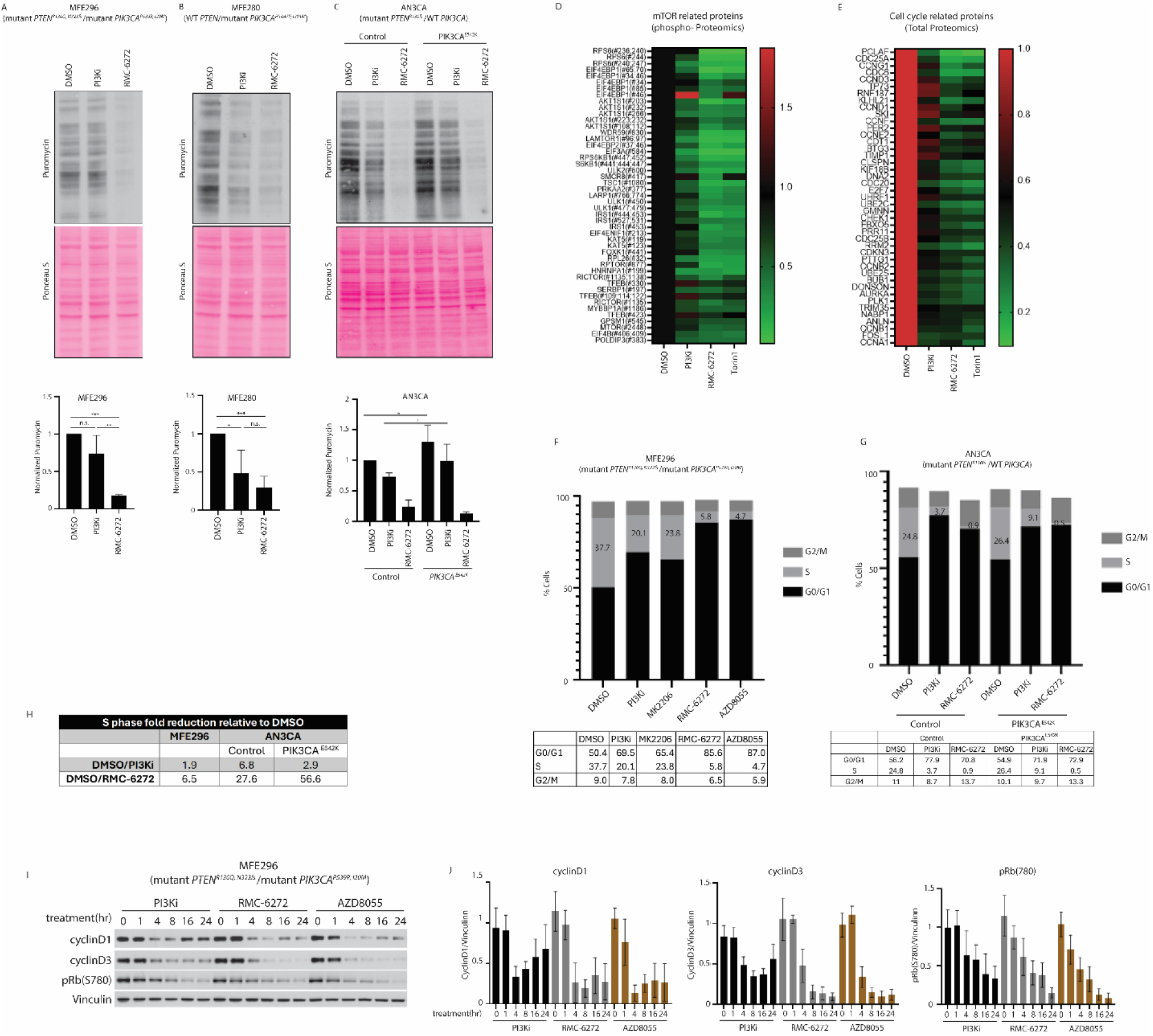
mTORC1 inhibition attenuates protein synthesis and cell cycle progression in *PTEN/PIK3CA* double mutant EC cells. MFE296 (mutant *PTEN^R130Q, N323fs^ /*mutant *PIK3CA^P539R, I20M^*) cells (A) and MFE280 (WT *PTEN/*mutant *PIK3CA^H1047Y; I391M^*) (B) were treated with a combination of 1μM BYL-719 + 250 nM AZD8186 (PI3Ki) or with 500 pM RMC-6272 for 24 hours followed by 30 minutes Puromycin (1μM) treatment. (C) AN3CA (mutant *PTEN^R130fs^ /*WT *PIK3CA*) cells were transiently transfected with mutant *PIK3CA^E542K^*, and after 24 hours were treated with either combination of 1μM BYL-719 and 250 nM AZD8186 (PI3Ki), or with 500pM RMC-6272 for additional 24 hours, followed by 30 minutes Puromycin (1μM) treatment. Western blots probed for puromycin show its incorporation into newly synthesized proteins (upper panel). Ponceau S labeling was used as a loading control (middle panel). Puromycin band intensity was quantified and normalized to Ponceau S and to DMSO treated control cells (lower panels). Graphs represent average and standard deviation of at least three independent experiments. ***p-value <0.001, **p-value <0.01, *p-value <0.05, n.s. not significant. (D-E) MFE296 (mutant *PTEN^R130Q, N323fs^ /*mutant *PIK3CA^P539R, I20M^*) cells were treated with either a combination of 1μM BYL-719 + 250 nM AZD8186 (PI3Ki), 500pM RMC-6272, or 250 nM Torin1 for 16 hours. Heat maps, displaying TMTpro-based quantification of indicated protein’s abundance, show the phosphorylated levels of mTOR-related proteins (D) and total expression of cell cycle regulators (E) upon treatment. (F-H) Cell cycle states of the following cells were analyzed by flow cytometry after inhibitors treatment followed by 2 hours EdU (10 μM) incorporation and EdU-FxCycle Violet staining. (F) MFE296 (mutant *PTEN^R130Q, N323fs^ /*mutant *PIK3CA^P539R, I20M^*) cells after treatment with a combination of 1μM BYL-719 + 250 nM AZD8186 (PI3Ki), 1μM MK2206, 500pM RMC-6272 or 500nM AZD8055 for 24 hours. (G) AN3CA (mutant *PTEN^R130fs^ /*WT *PIK3CA*) cells that were transiently transfected with mutant *PIK3CA^E542K^*, and after 24 hours were treated with either combination of 1μM BYL-719 and 250 nM AZD8186 (PI3Ki), or with RMC-6272 (500pM) for additional 24 hours. (H) A summarizing table of fold change in S phase relative to DMSO observed in F-G. Numbers represent an average of three independent experiments. (I-J) MFE296 (mutant *PTEN^R130Q, N323fs^ /*mutant *PIK3CA^P539R, I20M^*) cells were treated with a combination of 1μM BYL-719 and 250 nM AZD8186 (PI3Ki), 500pM RMC-6272, or 500nM AZD8055 for the indicated time points. The expression of the indicated cell cycle regulators was analyzed by immunoblotting using specific antibodies. Lysates from the experiment shown in Figure 3D were used for this analysis. (J) Bands were quantified and normalized to vinculin expression. Quantification demonstrates an average and standard deviation of 3-4 independent experiments.

These findings suggest that coexistent *PTEN* and *PIK3CA* alterations increase basal levels of mTORC1 signaling and cap-dependent translation. PI3K inhibition was unable to effectively inhibit the increased mTOR-driven protein synthesis in EC with coexistent *PTEN* loss and an activating *PIK3CA* mutation, while the mTORC1-selective inhibitor, RMC-6272, potently inhibited mTORC1 and protein translation in these EC cells.

To identify proteins responsible for the effects of mTOR inhibition on proliferation, an unbiased global proteomic analysis was performed. This was done with the multiplexed tandem mass tagging (TMT)pro-mass spectrometry. The double mutant MFE296 (mutant *PTEN^R130Q,N323fs^ /* mutant *PIK3CA^P539R,I20M^*) cells were treated for 16 hours with PI3K inhibitors, RMC-6272, or the dual mTORC1/2 inhibitor, Torin1, and changes in proteome and phospho-proteome were analyzed. We quantified the changes in expression and phosphorylation of proteins that were significantly altered by inhibitor treatment and analyzed their functional ontology. RMC-6272 reduced phosphorylation of 366 proteins, with mTOR signaling and protein translation emerging as the most prominently enriched processes. Fig. 4D shows a group of 47 mTOR-related proteins (e.g., 4E-BP1, S6K, S6, ULK1) with decreased phosphorylation following RMC-6272 or Torin1 treatment, whereas PI3K inhibition produced a much weaker effect, consistent with the data shown in Fig. 3D. RMC-6272 treatment also reduced the total expression of 140 proteins, with the most significantly affected group comprising of cell cycle regulators (e.g., CCND1, CCNE2, CCNB1), phosphatases (CDC25A/B), DNA replication and mitotic regulators (e.g., CDC6, AURKA, PLK1). Similar reductions were seen with Torin1 and with the PI3K inhibitors, but the latter led to a lesser decrease in these target proteins (Fig. 4E). Other ontology categories enriched by the treatments are presented in Supplementary Fig. 7. Of note, the increased expression of IRS1 is consistent with the known mTOR/S6K-mediated negative feedback loop ^33^.

### The PI3K/mTOR pathway drives G1 progression in EC tumors

EC tumors with concurrent *PIK3CA* and *PTEN* alterations are resistant to PI3K inhibition but remain sensitive to direct mTORC1 blockade, which effectively reduces cell growth, mTORC1-dependent translation, and cell-cycle regulator proteins. We therefore examined changes in the cell cycle in EC cell models. Treatment of MFE296 with either mTOR inhibitor, RMC-6272 or AZD8055 for 24 hours, substantially decreased S phase cells from 37.7% to 5.8% and 4.7% (respectively). G0/G1 phase increased from 50.4% to 85.6% and 87% (respectively) and cell growth was profoundly inhibited (Fig. 3C). In the cells treated with PI3K or AKT inhibitors for 24 hours, S-phase decreased with PI3K inhibitors (20.1%) or with AKT inhibitor MK2206 (23.8%), and G0/G1 phase increased to 69.5% and 65.4% (respectively) (Fig. 4F and Supplementary Fig. 8), and treatment of these cells with PI3K inhibitors slowed but did not arrest growth (Fig. 1D, 2B).

Treatment of the isogenic double mutant AN3CA cells with PI3K inhibitors, led to the reduction in quantity of cells in the S phase but less than their single mutant counterpart (Fig. 4G and Supplementary Fig. 9). This is consistent with the continued growth of double mutant cells on PI3K inhibitors while the single mutants grew significantly slower (Fig. 1F). By contrast, treatment with the mTORC1 inhibitor RMC-6272 halted cell growth, leaving fewer than 1% of cells in S phase in the double mutant cells (Fig. 4G and Supplementary Fig. 9). Therefore, PI3K inhibitors did not effectively inhibit cell cycle progression and cell growth in double mutant cells, whereas RMC-6272 effectively and significantly inhibited cell cycle progression and growth. Both PI3K inhibitors and RMC-6272 modestly increased cell death in double-mutant MFE296 cells with more cell death induced by PI3K inhibitors than RMC-6272 (Supplementary Fig. 10), suggesting that the primary antiproliferative effect of RMC-6272 in the EC cells was mediated by inhibition of cell cycle progression.

We investigated if the effect of mTORC1 inhibition on cell cycle progression and cell growth is due to inhibition of D-cyclins expression and attendant RB hypo-phosphorylation in the double mutant cells. Fig. 4I-J shows that mTOR kinase inhibitors caused deeper and more durable reduction in the expression of cyclin D1, cyclin D3 in the double mutant MFE296 cells as compared to PI3K inhibitors, which showed a greater rebound in protein levels at 16 or 24 hours. This was associated with decreased phosphorylation of Rb at residue S780 upon both treatments, with a more substantial reduction upon treatment with the mTOR kinase inhibitors after 16 hours. The pattern of decreased expression of cell cycle–related proteins upon mTOR or PI3K inhibition is consistent with the sensitivity of MFE296 cells to these treatments.

Overall, these data suggest that the effects of PI3K/AKT/mTOR pathway inhibitors on EC cell proliferation are primarily driven by cell cycle regulation. The *PTEN/PIK3CA* double mutant cells have high mTOR activity, therefore can sustain protein translation and maintain cyclin D1 levels even in the presence of PI3K or AKT inhibition. This enables them to progress into S phase and allows cell growth. However, direct inhibition of mTORC1 significantly inhibits protein translation, cyclin D1 and D3 expression and arrests double mutant cells in G0/G1 phase.

### The selective bi-steric mTORC1 inhibitor, RMC-6272, suppresses the growth of *PTEN/PIK3CA* double mutant EC tumors *in vivo*

We assessed the sensitivity of the MFE296 (mutant *PTEN^R130Q,N323fs^ /* mutant *PIK3CA^P539R,I20M^*) cell-line derived xenograft model (CDX) to RMC-6272, AZD8055 or the combination of BYL719 and AZD8186 (PI3K inhibitors) in immunodeficient mice. Consistent with data from the *in vitro* experiments, treatment with PI3K inhibitors slowed but did not block tumor growth *in vivo*. By contrast, both mTOR kinase inhibitors, as single agents, completely inhibited tumor growth *in vivo* (Fig. 5A). The effect of RMC-6272 was comparable to that of AZD8055, suggesting that inhibition of mTORC1 activity may be sufficient to block the growth of MFE296-derived tumors. We found that 4E-BP1 and ULK1 phosphorylation levels were reduced after 4 hours of RMC-6272 or AZD8055 treatment *in vivo* and remained low for 24 hours (∼3.5 and ∼3.8-fold decrease in average), while their levels were less affected upon PI3K inhibition (∼1.5-fold decrease). Cyclin D1 and cyclin D3 levels were reduced durably after AZD8055 or RMC-6272 treatment (∼2-fold decrease in average) but not by PI3K inhibitors (Fig. 5B-C). The observed pathway modulation of mTOR inhibitors is consistent with their sustained inhibition on tumor growth, whereas PI3K inhibition produced only a marginal effect in the double mutant model.

**Figure 5:**
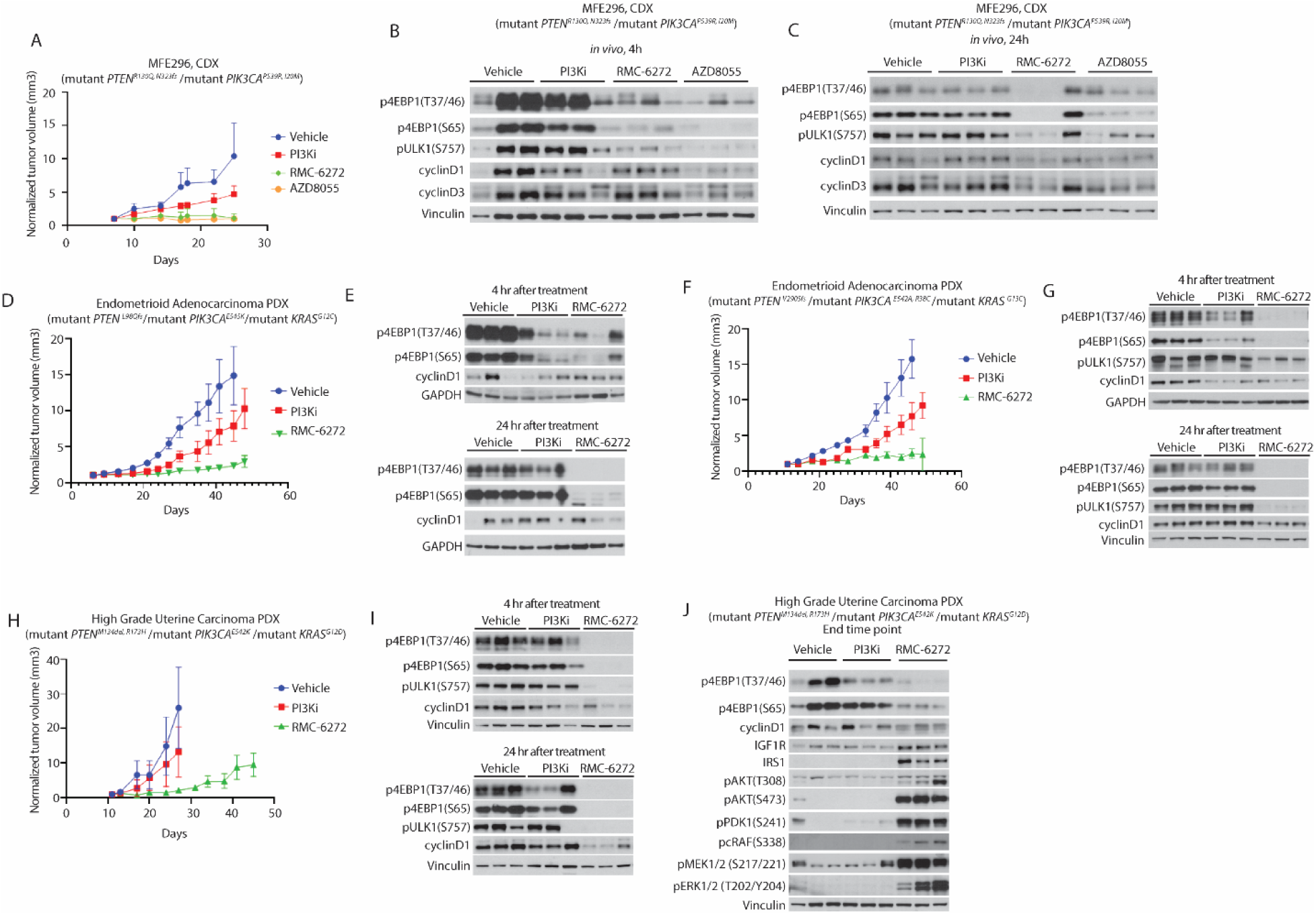
RMC-6272 suppresses the growth of *PTEN/PIK3CA* double mutant EC tumors *in vivo*. (A-C) Mice bearing MFE296 (mutant *PTEN^R130Q, N323fs^ /*mutant *PIK3CA^P539R, I20M^*) xenograft (CDX) tumors were treated with either combination of BYL719 (25 mg/Kg p.o QDx5) and AZD8186 (75 mg/Kg p.o. BIDx5) (PI3Ki), RMC-6272 (3 mg/kg i.p. QW), or AZD8055 (75 mg/Kg p.o. 3 times/week). Tumor volumes were measured and mean, and SD values presented (n = 5). (B-C) Immunoblots depict mTOR output after 4 hours (B) or 24 hours (C) *in vivo* treatment (n = 3). (D-J) Mice bearing the indicated EC PDX xenograft models were treated with either combination of BYL719 (25 mg/Kg p.o QDx5) and AZD8186 (75 mg/Kg p.o. BIDx5) (PI3Ki) or RMC-6272 (3 mg/kg i.p. QW). (D-E) Endometrioid adenocarcinoma (mutant *PTEN ^L98Qfs^/*mutant *PIK3CA ^E545K^/*mutant *KRAS ^G12C^*). (F-G) Endometrioid adenocarcinoma (mutant *PTEN ^V290Sfs^ /*mutant *PIK3CA ^E542A, R38C^ /*mutant *KRAS ^G13C^*). (H-J) High-grade uterine carcinoma (mutant *PTEN ^M134del, R173H^ /* mutant *PIK3CA ^E542K^ /*mutant *KRAS ^G12D^*). (D, F, H) Tumor volumes were measured and mean, and SD values represented (n = 5). (E,G,I) mTOR output assessed by immunoblotting after 4 hours treatment (upper panels) or 24 hours treatment (lower panels) (n = 3). (J) Immunoblots depicting PI3K/mTOR signaling output in 3 representative high-grade uterine carcinoma (mutant *PTEN ^M134del, R173H^/*mutant *PIK3CA ^E542K^ /*mutant *KRAS ^G12D^*) tumors collected at the end of the experiment shown in (H).

We further tested additional EC derived PDX models containing *PTEN/PIK3CA* double mutations. These include endometrioid adenocarcinoma (mutant *PTEN^L98Qfs^/*mutant *PIK3CA^E545K^/*mutant *KRAS^G12C^*), endometrioid adenocarcinoma (mutant *PTEN^V290Sfs^ /* mutant *PIK3CA^E542A, R38C^ /*mutant *KRAS^G13C^*), and high-grade uterine carcinoma (mutant *PTEN^M134del,R173H^ /*mutant *PIK3CA^E542K^ /*mutant *KRAS^G12D^*). Approximately 30% of endometrial tumors with coexistent *PTEN* and *PIK3CA* lesions also harbor *KRAS* mutations (Supplementary Fig. 1C-D). The growth of these tumors was moderately controlled by PI3K inhibitors. On the other hand, mTORC1 inhibition by RMC-6272 caused significant suppression of tumor growth *in vivo* for approximately 40 days in two of the models (Fig. 5D, F, H). The high-grade uterine carcinoma PDX (mutant *PTEN^M134del, R173H^ /*mutant *PIK3CA^E542K^ /*mutant *KRAS^G12D^*) underwent growth arrest with RMC-6272 treatment for 16 days, followed by moderate re-growth of the tumors (Fig. 5H). In all models, phosphorylation of mTORC1 targets (4E-BP1, ULK1) was reduced 4 hours after RMC-6272 treatment by ∼70-fold vs. vehicle and remained low for 24 hours, while the reduction in cyclin D1 expression was prominent at 24 hours after treatment (∼2-fold vs. vehicle in average). mTORC1 targets (4E-BP1, ULK1) phosphorylation and cyclinD1 levels were less affected by PI3K inhibition with average of 1.5-fold reduction compared with vehicle (Fig. 5E, G, I). Furthermore, RMC-6272 did not induce hyperglycemia or cause weight loss in mice (Supplementary Fig. 11).

### Combined inhibition of mTORC1 and mutant KRAS signaling leads to improved anti-tumor activity in *PTEN/PIK3CA* double mutant EC tumors with co-occurring *KRAS* mutation

KRAS mutations occur in 18-21% of patients with endometrial cancer, but its occurrence is enriched in endometrial cancer harboring *PTEN* and *PIK3CA* alterations, reaching 25-33% (Supplementary Fig. 1). The high-grade uterine carcinoma PDX model (mutant *PTEN^M134del, R173H^ /* mutant *PIK3CA^E542K^ /mutant KRAS^G12D^*) that relapsed on RMC-6272 treatment (Fig. 5H) was examined in more detail. We found that while phosphorylation of 4E-BP1 remained low with RMC-6272 treatment, there was an increase in phosphorylation of cRAF, MEK, ERK, PDK1 and AKT (Fig. 5J). The expression of IGF1R and IRS1 was elevated compared to that of tumors treated with vehicle or PI3K inhibitors likely due to relief of feedback inhibition of RAS signaling and/or RTK signaling ^33^.

Since this high-grade uterine carcinoma PDX model harbors the KRAS^G12D^ mutation, we next asked whether RAS activation played a role to support tumor cell survival and re-growth and whether combined mTORC1 and RAS inhibition could prevent tumor regrowth.

We treated the high-grade uterine carcinoma (mutant *PTEN^M134del, R173H^ /*mutant *PIK3CA^E542K^ /mutant KRAS^G12D^*) PDX tumor-derived cells *in vitro* with RMC-6272 (500 pM) together with the RAS(ON) multi-selective inhibitor, RMC-7977 (100 nM). RMC-7977 is a preclinical tool compound representative of daraxonrasib (RMC-6236), and both compounds are structurally related and have comparable *in vitro* and *in vivo* properties in preclinical models ^34^. *In vitro* cell growth was inhibited by RMC-7977 and RMC-6272 individually, with RMC-6272 showing a more pronounced effect. The combination treatment resulted in a slightly better attenuation of cell growth (Supplementary Fig. 12, upper panel). RMC-6272 treatment briefly induced phosphorylation of RAS/MAPK pathway markers c-RAF, MEK and ERK at about 4-hour (Supplementary fig. 12 lower panel). RMC-7977, either alone or in combination with RMC-6272 inhibited the phosphorylation of these RAS/MAPK pathway markers and reduced DUSP6 levels. Phosphorylation of mTORC1 targets (4E-BP1, ULK1, S6K, and S6) was minimally affected by RAS inhibition but suppressed by RMC-6272 alone or in combination with RMC-7977. Interestingly, in comparison to single agent treatment, the combination of RMC-6272 with RMC-7977 further reduced the phosphorylation of AKT, its substrates FOXO and PRAS40, and the expression of cyclin D1 and cyclin D3 along with increases in apoptosis markers cleaved PARP and cleaved caspase-3 (Supplementary Fig. 12 lower panel). Similar combinatorial effects were noted in three other EC models with coexistent *PTEN/PIK3CA* double mutations and mutant *KRAS.* These results were consistent with those we observed in KRAS G12C lung cancer models treated with the combination of RAS and mTORC1 inhibitors ^35^ (Supplementary Fig. 13-15).

To test the effects of combination *in vivo*, we evaluated anti-tumor activity of RMC-6272, RMC-7977, and their combination in mice bearing these human EC PDX models: (1) high-grade uterine carcinoma (mutant *PTEN^M134del, R173H^ /*mutant *PIK3CA^E542K^ /*mutant *KRAS ^G12D^*), (2) endometrioid adenocarcinoma (mutant *PTEN^V290Sfs^ /*mutant *PIK3CA^E542A, R38C^ /*mutant *KRAS^G13C^*), and (3) endometrioid adenocarcinoma (mutant *PTEN ^L98Qfs^/* mutant *PIK3CA^E545K^/* mutant *KRAS^G12C^*). The latter was also treated with the RAS(ON) G12C-selective inhibitor, RMC-4998, which is a preclinical tool compound representative of the investigational agent elironrasib (RMC-6291) ^36^. In all three models, RAS inhibition alone was ineffective on tumor growth *in vivo*, however, the combination of RMC-6272 and RAS(ON) inhibitor delayed the onset of treatment-resistant tumor growth that was observed with RMC-6272 alone (Fig. 6A, C and E). In general, RMC-6272 inhibited the phosphorylation of mTORC1 targets 4E-BP1, S6K, and ULK1 at 4 and 24 hours after treatment, with the exception of the KRAS^G12D^ high-grade uterine carcinoma model that showed p4E-BP1 reduction at 24 hours. RMC-7977 did not cause significant effects on phosphorylation of these mTORC1 targets, except for ULK1 phosphorylation in the endometrioid adenocarcinoma (mutant *PTEN^V290Sfs^ /*mutant *PIK3CA^E542A, R38C^ /*mutant *KRAS^G13C^*) (Fig 6D). ERK phosphorylation was induced upon RMC-6272 treatment in the endometrioid adenocarcinoma model (mutant *PTEN^V290Sfs^ /*mutant *PIK3CA^E542A, R38C^/*mutant *KRAS^G13C^*), but it was completely abolished by RMC-7977. Overall, the combination treatment led to further reduction of DUSP6 and cyclin D1 levels in all three models as compared to single agent mTORC1 inhibition alone. Cyclin D3 levels were mostly affected by RMC-6272 alone and in the combination treatment. Although either mTORC1 or RAS(ON) single agent treatment led to cleaved caspase-3 and PARP, the combination further induced these apoptosis markers, with the exception of one endometrioid adenocarcinoma (mutant *PTEN^L98Qfs^/* mutant *PIK3CA^E545K^/* mutant *KRAS^G12C^*) model that showed only a transient induction (Fig. 6B, D, F and Supplementary Fig. 16). Altogether, our data show that in *PTEN/PIK3CA/KRAS* triple mutant tumors, combined mTORC1 and KRAS inhibition suppressed both mTORC1 and RAS signaling outputs better than single agents alone, which resulted in a further reduction of cyclin D1 levels and importantly, an increase in cleaved caspase-3 and PARP levels. The combination treatment of RMC-6272 and RMC-7977 demonstrated improved anti-tumor activity *in vivo* as compared to either single agent in our preclinical models (Fig. 6A, C and E). Furthermore, the blood glucose concentrations in mice treated with the combination did not show a signal of hyperglycemia, and mice tolerated the treatment based on body weight assessment (supplementary Fig.11).

**Figure 6:**
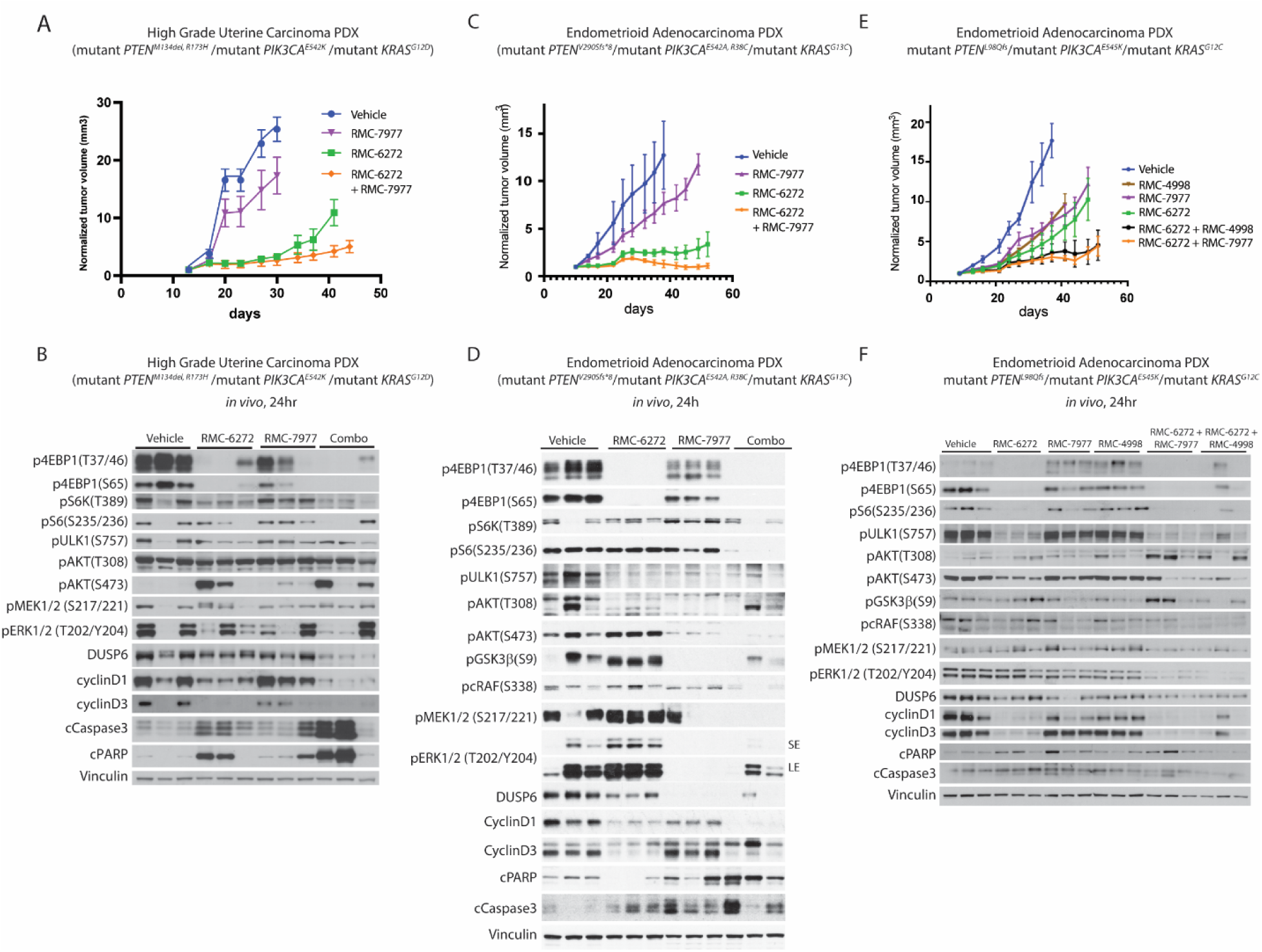
Combined inhibition of mTORC1 and KRAS improves anti-tumor activity in *PTEN/PIK3CA* double mutant EC tumors with co-occurring *KRAS* mutation. Mice bearing the high-grade uterine carcinoma (mutant *PTEN ^M134del, R173H^ /*mutant *PIK3CA ^E542K^ /*mutant *KRAS ^G12D^*) PDX were treated *in vivo* with RMC-6272 (3 mg/kg i.p. QW), RMC-7977 (25mg/kg p.o. 3 day/week) or their combination. Tumor volumes were measured and mean, and SD values presented (n = 5). (B) Immunoblots depict RAS/PI3K/mTOR signaling output after 24 hours of *in vivo* treatment. (C) Endometrioid adenocarcinoma (mutan*t PTEN ^V290Sfs^ /*mutant *PIK3CA ^E542A, R38C^ /*mutant *KRAS ^G13C^*) PDX cells were cultured *in vitro* and subsequently implanted into mice to establish cell-derived xenograft (CDX) tumors. Mice were treated *in vivo* with RMC-6272 (3 mg/kg i.p. QW), RMC-7977 (25mg/kg p.o. 3 day/week) or their combination. Tumor volumes were measured and mean, and SD values presented (n = 6). (D) Immunoblots depict RAS/PI3K/mTOR signaling output after 24 hours of *in vivo* treatment. Notes: pERK was examined with short exposure (SE) and long exposure (LE). (E) Endometrioid adenocarcinoma (mutant *PTEN ^L98Qfs^/*mutant *PIK3CA ^E545K^/*mutant *KRAS ^G12C^*) PDX cells were cultured *in vitro* and subsequently implanted into mice to establish cell-derived xenograft (CDX) tumors. Mice were treated *in vivo* with RMC-6272 (3 mg/kg i.p. QW), RMC-7977 (25mg/kg p.o. 3 day/week), RMC-4998 (80mg/kg p.o. QDx5) or either RMC-6272+ RMC-7977 or RMC-6272+ RMC-4998 combinations. Tumor volumes were measured and mean, and SD values presented (n = 6). (F) Immunoblots depict RAS/PI3K/mTOR signaling output after 24 hours of *in vivo* treatment.

## Discussion

The high frequency of coexistent inactivating *PTEN* and activating *PIK3CA* mutations in endometrial cancer is unique. While we do not understand the predilection for coexistence of these mutations in EC tumors, their prevalence suggests that activation of PI3K/mTOR signaling plays an important role in the biology of this disease. However, clinical studies of rapamycin-like drugs in endometrial cancers revealed only modest clinical benefit and did not support this hypothesis ^11,13–16^.

In previous work, we discovered that PTEN translation is controlled by PI3K/mTOR, such that PI3K activation enhances PTEN expression whereas PI3K pathway inhibition reduces it ^6^. Since PTEN is a phosphatase that negatively regulates the PI3K pathway by dephosphorylating PIP3 the lipid product of PI3K signaling, changes in PTEN expression buffer changes in the output of the PI3K/mTOR pathway. Elevated PI3K signaling increases PTEN expression which in turn downregulates pathway output. Thus, PTEN acts as a downstream feedback regulator of the PI3K/AKT/mTOR pathway. Similarly, activated mTOR or AKT reduces upstream signaling by inhibiting the expression and function of some activated receptor tyrosine kinases ^5,33,37^. We previously hypothesized that in tumors in which PI3K is mutated and PTEN function is lost, both upstream and downstream feedback inhibition are impaired so that pathway output would be hyperactivated. This turned out to be the case and, in such models, basal mTORC1 kinase signaling is remarkably elevated and cooperates with PI3K signaling to hyperactivate tumor cell migration and invasion ^6^.

Now, in this study we show that coexistent PI3K mutation and loss of PTEN phosphatase promoted elevated levels of PIP3 relative to either alteration alone with corresponding increases in mTOR kinase activity and mTOR driven cap-dependent translation in EC. In these double mutant models, PI3K or AKT inhibitors were insufficient to reduce mTORC1 signaling, protein translation and cell growth. Knockout of *PTEN* in a *PIK3CA* mutant cell led to a significant increase in PIP3, 4E-BP1 and ULK1 phosphorylation while overexpression of *PIK3CA* mutation in a *PTEN* mutant cell led to increased activation of pAKT and pS6K.

While pan-mTOR catalytic inhibitors could be used to block mTOR activity in EC, the clinical utility of these agents has been limited by their toxicity, including induction of hyperglycemia due to inhibition of phosphorylation of AKT S473 by mTORC2 kinase^22,38^. The findings presented here demonstrate that mTORC1 selective inhibitors may offer an alternative approach for the management of endometrial cancers. RMC-6272 is such a bi-steric mTORC1-selective inhibitor that has antitumor activity *in vivo* and does not dephosphorylate AKT S473 nor cause hyperglycemia^28,38^. In contrast to rapalog treatment and/or PI3K inhibition, RMC-6272 profoundly inhibited mTORC1 and its downstream sequelae in *PIK3CA/PTEN* double mutant EC cells, as evidenced by reduced 4E-BP1 phosphorylation, inhibition of cap-dependent translation and a reduction in cyclin D1 levels, the number of cells in S phase, and cell proliferation *in vitro*. RMC-6272 also inhibited growth of endometrial tumor CDXs and PDXs *in vivo*, resulting in tumor regression in some cases.

The related bi-steric mTORC1-selective inhibitor RMC-5552 was evaluated in a single arm phase 1/1b dose escalation study with 57 patients, 4 of whom had endometrial cancer refractory to other therapies. Three of these patients had coincident *PTEN* inactivating and *PIK3CA* activating mutations, of which one had a complete response for over 6 months as of the June 2024 data cut. Furthermore, a patient with a salivary gland tumor harboring a *PTEN* mutation demonstrated a partial response, with 78% tumor reduction. This observation suggests the potential relevance of RMC-5552 treatment for tumors with *PTEN* mutations in addition to those with *PTEN/PIK3CA* double mutations. In this single arm study, hyperglycemia, which is frequently observed with mTORC1/2 kinase inhibitor was infrequent (4%) and not dose-limiting, consistent with the mTORC1 selectivity of RMC-5552 at clinically active doses ^39^.

While these findings are encouraging much remains to be learned about the biology of endometrioid tumors and how to treat them. In the PDX models, whereas treatment with RMC-6272 for up to two months arrested tumor growth there was negligible induction of cell death and no measurable regression. Endometrioid tumors have additional somatic mutations, including in *KRAS*, which is a known oncogenic driver and 25-33% of patients with *PTEN* and/or *PIK3CA* mutations also carry mutant *KRAS* (Supplementary Fig. 1). We found that all the PDXs we examined had *KRAS* hotspot mutations (Fig 6). Previously we have shown that combined inhibition of KRAS^G12C^ and mTORC1 resulted in synergistic inhibition of proliferation and induction of apoptosis in preclinical models of KRAS^G12C^ non-small cell lung cancer ^28,35^. Here, we demonstrate that co-targeting KRAS and mTORC1 improved anti-tumor activity in triple mutant EC models harboring alterations in *KRAS*, *PTEN* and *PIK3CA*. The combined inhibition of KRAS and mTORC1 resulted in a stronger reduction of mTORC1 and RAS signaling output and cyclin D1 expression which translated into improved antitumor activity, although broad tumor regressions were still not observed.

Thus, our preclinical studies have shown that a consequence of the co-occurrence of *PIK3CA* mutations and *PTEN* loss in endometrial cancer is hyperactivation of mTORC1 kinase signaling that leads to increased translation of cell cycle regulators like cyclin D1 and increased cell growth. PI3K inhibitors or rapalogs are insufficient to inhibit mTORC1 kinase in tumors with both mutations, but mTORC1-selective kinase inhibitors are, and cause growth arrest of the tumors without causing hyperglycemia. These preclinical results support the clinical evaluation of investigational agents such as RMC-5552 in EC with coexistent *PIK3CA* mutation and *PTEN* loss, along with combination therapies such as RAS(ON) inhibitors to enhance antitumor activity.

## Methods

### Cell culture and reagents

The endometrial cancer cell lines KLE, MFE280, AN3CA, MFE296, NOU1, HEC6 and the endometrial carcinoma PDX-derived cells were grown in DMEM-F12 medium supplemented with 2mM glutamine, 50 U/mL penicillin, 50 μg/mL streptomycin, and 10% fetal bovine serum. All cells were maintained at 37°C in 5% CO2. BYL719, rapamycin, MK2206 were obtained from Selleck Chemical. AZD8186, AZD8055 and AZD5363 were provided by AstraZeneca. RMC-6272, RMC-7977, and RMC-4998 were provided by Revolution Medicines. Torin1 was obtained from Cell Signaling Technology (#14379). Compounds were dissolved in DMSO to a final concentration of 10 mmol/l and stored at –20°C. Puromycin was obtained from Gibco (Thermo Fisher Scientific).

### Immunoblotting

Cells in culture were washed in cold PBS and collected to pellets, followed by lysis with Cell Lysis Buffer (Cell Signaling #9803) supplemented with Halt protease and phosphatase inhibitors (Pierce Chemical). Lysates were briefly sonicated before centrifugation at 20,817g for 15 minutes at 4°C.

Xenograft tumors were homogenized in SDS lysis buffer (50mM Tris-HCL pH 7.4, 10% Glycerol, 2% SDS) and boiled at 100°C for five minutes. Lysates were then briefly sonicated, boiled again for 5 minutes, before clearing by centrifugation at 20,817g for 10 minutes at room temperature.

The supernatant was collected, and protein concentration was determined using the BCA kit (Pierce) according to the manufacturer’s instructions. Protein samples were diluted in 4X LDS sample Buffer with 10X Sample Reducing Agent (both from Invitrogen). 20 μg of protein was loaded onto each lane of a 4%–12% BisTris mini gel or midi gel (Invitrogen) for immunoblotting. Transfer was onto nitrocellulose membranes (0.2 mm, GE Health Care) before blocking for 1h at room temperature and incubating with primary antibodies overnight at 4°C. Membranes were incubated with secondary rabbit antibody (Sigma) or secondary mouse antibody (GE Health Care) for 1h at room temperature. Blots were developed in Amersham ECL detection reagent (Cytiva) or Millipore’s Immobilon HRP reagents according to the manufacturer’s instructions. Bands quantification was conducted by ImageJ software.

Primary antibodies obtained from Cell Signaling Technologies and used at 1:1000 dilution: pAKT T308 (#2965), pAKT S473 (#4060), AKT (#9272), pGSK3β S9 (#9323), pPRAS40 T246 (#2997), p4EBP1 T37/46 (#9459), p4EBP1 S65 (#9451), pULK1 S757(#6888), p-p70S6K T389 (#9234), pS6 S235/236 (#4858), vinculin (#13901), PTEN (#9559), Histone H3 (#4499), Flag (98533), cyclinD1 (#55506), cyclin D3 (#2936), pRb S780 (#9307), pcRAF S338 (#9427), pMEK S217/221 (#9154), pERK T02/Y204 (#4370), ERK (#4696), pPDK1(S241) (#3438), IGF1R (#3027), IRS1 (#3407), Phospho-FoxO1 (T24)/FoxO3a (T32) (#9464), cleaved PARP (#5625), cleaved Caspase-3 (#9661), , 4EBP1 (#9452), Actin (#4970).

Other antibodies: DUSP6 (Abcam, ab76310), Puromycin (Kerafast, EQ0001)

All Western blot experiments were repeated at least twice, and a representative result is shown.

### Cell growth assay

Cells were plated at 2000 cells per well in a 96-well plate and grown in 8 replicates per condition, then treated with inhibitor the following day. At indicated times, plates were treated with either alamarBlue (DAL1025, Thermo Fisher Scientific), or with PrestoBlue (A13262, Thermo Fisher Scientific) and assay was conducted according to manufacturer instructions. All cell growth experiments were repeated at least three times, and a representative result is shown.

MFE280 cells (200,000) in Figures 1E and 2D were plated on 6 well plates. At indicated times, cells were removed by trypsin and directly counted using trypan blue staining and hemocytometer.

### Transfections and retroviral infections and plasmids

For mutant *PIK3CA*^E542K^ over-expression, we first introduced the mutation to the pLP-LNCX-*PIK3CA*-WT plasmid using the site-directed mutagenesis Kit (QuikChange, #200521, Agilent) using the following primers:

F: ACA CGA GAT CCT CTC TCT AAA ATC ACT GAG CAG GAG AAA

R: TTT CTC CTG CTC AGT GAT TTT AGA GAG AGG ATC TCG TGT

pLP-LNCX-*PIK3CA*-WT was a gift from Todd Waldman (Addgene plasmid #25633)

pLNCX2 Retroviral Vector was obtained from Takara (#631503) and served as a control for transfections.

GFP-PH^AKT^ plasmid was described ^22^

Transient transfections were conducted using the Lipofectamine 3000 Transfection Reagent (Thermo Fisher Scientific, L3000001)

Lenti viral infections were carried out as previously described ^38^.

pLKO-PTEN-shRNA-3001 was a gift from Todd Waldman (Addgene plasmid #25639). The nontargeting control hairpin (shControl) used is #SHC016. Cells were selected using puromycin (5 μg/ml)

### EdU-DAPI based flow cytometry

3×10^6^ cells were plated in 10 cm dishes and treated with the indicated inhibitors for 24 hours. Before harvesting, cells were incubated with 10 μM EdU for two hours. The cells were harvested stained with Click-iT EdU Alexa Fluor 488 Flow Cytometry Assay Kit (Invitrogen) according to the manufacturer’s protocol. Data were obtained on an LSR-II analyzer and analyzed with DIVA software.

### Annexin V-propidium iodide (PI) assay

5–10 × 10^5^ cells were plated in 10 cm dishes and treated with the indicated inhibitors After 72 h, floating cells in media and adherent, trypsinized cells were collected in a single tube and stained with annexin V and propidium iodide using the FITC annexin V Apoptosis Detection Kit I (BD Biosciences) according to the manufacturer’s protocol. Data were obtained on a LSRFortessa flow cytometer and analyzed with Diva software.

### Xenograft studies

Tumor specimens from uterine endometrioid carcinoma patients were collected under an approved IRB protocol (protocol #14-091). Tumor tissue was immediately minced, mixed (50:50) with matrigel (Corning, New York, NY) and implanted subcutaneously in 6-8 weeks old female NSG mice (Jackson Laboratory, Bar Harbor, ME) to generate Patient Derived Xenografts (PDX) as previously described ^40^. Mice were monitored daily, and models were transplanted in mice three times before being deemed established. PDX tumor histology was then confirmed by pathology review of H&E slides, and direct comparison to the corresponding patient slides.

For efficacy studies, established PDXs were serially transplanted into 6-8 week old female NSG mice as above. Once tumors reached an average volume of 100-150 mm^3^, mice were randomized to receive either a vehicle control, BYL719 (25 mg/Kg p.o QDx5) + AZD8186 (75 mg/Kg p.o. BIDx5), RMC-6272 (3 mg/kg i.p. QW), RMC-7977 (25mg/kg p.o. 3 day/week), RMC-4998 (80mg/kg p.o. QDx5) or their combinations as indicated in the figures. NSG mice s.c. engrafted with 10 million MFE296 cells, were similarly randomized when tumors reached an average volume of 100-150 mm^3^ and treated with the same compound combinations.

In all instances, mice were observed daily throughout the treatment period for signs of morbidity/mortality. Tumors were measured twice weekly using calipers, and tumor volume was calculated using the formula: length x width^2^ × 0.52. Body weight was also assessed twice weekly. After ∼4 weeks of treatment tumor samples were collected for further analysis.

Animal experiments performed at Memorial Sloan Kettering (MSK) were done according to the protocol approved by the MSK Animal Care and Use Committee.

### PIP3 detection and ImageStreamX analysis

Cells were transfected with a plasmid expressing GFP fused to the PH domain of AKT (GFP-PH^AKT^), as described ^22^. ImageStream^X^, an imaging flow cytometer was used to analyze the fluorescent pattern in each cell in flow. Using the masking tool, the median GFP intensity in the plasma membrane and cytoplasm was determined for each cell. The ratio of these two measurements was then calculated (PM/cytoplasm). This ratio was plotted in histograms, showing the frequency of cells in the population relative to their score. Using the Julius tool, statistical significance was calculated by pairwise Welch t-test comparing the group means, including 95% CIs and Benjamini–Hochberg FDR adjustment.

### Mass spectrometry

Unbiased global proteomic analysis was performed with the multiplexed tandem mass tagging TMT (Tandem Mass Tag) pro-mass spectrometry. The double mutant MFE296 (mutant *PTEN^R130Q,N323fs^* /mutant *PIK3CA^P539R,I20M^*) cells were treated for 16 hours with 1μM BYL-719 + 250 nM AZD8186 (PI3Ki), 500pM RMC-6272, or 250 nM Torin1 in triplicate, and proteome and phospho-proteome changes were analyzed following the method described in Yi et al ^41^. Protein quantification values were exported for further analysis in Microsoft Excel and Perseus. In Perseus, a two-way Welch’s t-test analysis was performed to compare two datasets, using the S0 parameter of 0.585 (a minimal fold change cutoff), and correction for multiple comparisons was achieved using the permutation-based FDR method, both of which are built-in functions of Perseus software. Each reporter ion channel was summed across all quantified proteins and normalized, assuming equal protein loading across all samples. The maximum and minimum TMT ratio quantifiable was capped at 100-fold. GO analysis was performed using Enrichr. Briefly, statistically significant protein hits were further filtered by p-value ≤0.01 and fold change cutoff of 2 (≥2-fold increase or ≤50% decrease). Enrichment analysis was performed on these protein sets using the ‘Gene Ontology Biological Process’ (Enrichr) as annotation source.

### Statistical analysis

The details of the statistical analysis of experiments can be found in the figure legends.

Statistical analyses were performed by an unpaired, 2-tailed Student t-test and P < 0.05 was defined as significant.

Independent experiments were conducted with a minimum of two biological replicates per condition to allow for statistical comparison.

Data are shown as mean ± SD.

## Acknowledgements

We thank Zannatul Monia, head of the flow cytometry core facility team at the New York blood center for her assistance with the ImageStream^X^ experiments. Funding: This research was supported by grants (to N.R.) from the National Institutes of Health (NIH) P01-CA129243; R35 CA210085; the Geoffrey Beene Cancer Research Center; the Emerson Collective Research Grant; Melanoma Research Alliance; The NIH MSKCC Cancer Center Core Grant P30 CA008748 and Experimental Therapeutics Center.

## Declaration of interests

N.R. is on the scientific advisory board (SAB) and owns equity in Beigene, Zai Labs, MapKure, Ribon and Effector. N.R. is also on the SAB of Astra Zeneca and Chugai and a past SAB member of Novartis, Millennium-Takeda, Kura, and Araxes. N.R. is a consultant to Revolution Medicines, Tarveda, Array-Pfizer, Boehringer-Ingelheim and Eli Lilly. He receives research funding from Revolution Medicines, AstraZeneca, Array, Pfizer and Boehringer-Ingelheim and owns equity in Kura Oncology and Fortress.

## References

1 Murali, R., Soslow, R. A. & Weigelt, B. Classification of endometrial carcinoma: more than two types. Lancet Oncol 15, e268–278 (2014). 10.1016/S1470-2045(13)70591-6

2 Paddock, M. N., Field, S. J. & Cantley, L. C. Treating cancer with phosphatidylinositol-3-kinase inhibitors: increasing efficacy and overcoming resistance. J Lipid Res 60, 747–752 (2019). 10.1194/jlr.S092130

3 Laplante, M. & Sabatini, D. M. mTOR signaling in growth control and disease. Cell 149, 274–293 (2012). 10.1016/j.cell.2012.03.017

4 Vasan, N. & Cantley, L. C. At a crossroads: how to translate the roles of PI3K in oncogenic and metabolic signalling into improvements in cancer therapy. Nat Rev Clin Oncol 19, 471–485 (2022). 10.1038/s41571-022-00633-1

5 Chandarlapaty, S. Negative feedback and adaptive resistance to the targeted therapy of cancer. Cancer Discov 2, 311–319 (2012). 10.1158/2159-8290.CD-12-0018

6 Mukherjee, R. et al. Regulation of PTEN translation by PI3K signaling maintains pathway homeostasis. Mol Cell 81, 708–723 e705 (2021). 10.1016/j.molcel.2021.01.033

7 Cerami, E. et al. The cBio cancer genomics portal: an open platform for exploring multidimensional cancer genomics data. Cancer Discov 2, 401–404 (2012). 10.1158/2159-8290.CD-12-0095

8 de Bruijn, I. et al. Analysis and Visualization of Longitudinal Genomic and Clinical Data from the AACR Project GENIE Biopharma Collaborative in cBioPortal. Cancer Res 83, 3861–3867 (2023). 10.1158/0008-5472.CAN-23-0816

9 Gao, J. et al. Integrative analysis of complex cancer genomics and clinical profiles using the cBioPortal. Sci Signal 6, pl1 (2013). 10.1126/scisignal.2004088

10 Alexa, M., Hasenburg, A. & Battista, M. J. The TCGA Molecular Classification of Endometrial Cancer and Its Possible Impact on Adjuvant Treatment Decisions. Cancers (Basel) 13 (2021). 10.3390/cancers13061478

11 Avila, M., Grinsfelder, M. O., Pham, M. & Westin, S. N. Targeting the PI3K Pathway in Gynecologic Malignancies. Curr Oncol Rep 24, 1669–1676 (2022). 10.1007/s11912-022-01326-9

12 Ibanez, K. R., Huang, T. T. & Lee, J. M. Combination Therapy Approach to Overcome the Resistance to PI3K Pathway Inhibitors in Gynecological Cancers. Cells 13 (2024). 10.3390/cells13121064

13 Saxton, R. A. & Sabatini, D. M. mTOR Signaling in Growth, Metabolism, and Disease. Cell 168, 960–976 (2017). 10.1016/j.cell.2017.02.004

14 Slomovitz, B. M. et al. A phase 2 study of the oral mammalian target of rapamycin inhibitor, everolimus, in patients with recurrent endometrial carcinoma. Cancer 116, 5415–5419 (2010). 10.1002/cncr.25515

15 Sun, S. Y. mTOR-targeted cancer therapy: great target but disappointing clinical outcomes, why? Front Med 15, 221–231 (2021). 10.1007/s11684-020-0812-7

16 Wang, Q., Peng, H., Qi, X., Wu, M. & Zhao, X. Targeted therapies in gynecological cancers: a comprehensive review of clinical evidence. Signal Transduct Target Ther 5, 137 (2020). 10.1038/s41392-020-0199-6

17 Weigelt, B., Warne, P. H., Lambros, M. B., Reis-Filho, J. S. & Downward, J. PI3K pathway dependencies in endometrioid endometrial cancer cell lines. Clin Cancer Res 19, 3533–3544 (2013). 10.1158/1078-0432.CCR-12-3815

18 Bonneau, D. & Longy, M. Mutations of the human PTEN gene. Hum Mutat 16, 109–122 (2000). 10.1002/1098-1004(200008)16:2<109::AID-HUMU3>3.0.CO;2-0

19 Han, S. Y. et al. Functional evaluation of PTEN missense mutations using in vitro phosphoinositide phosphatase assay. Cancer Res 60, 3147–3151 (2000).

20 Vasan, N. et al. Double PIK3CA mutations in cis increase oncogenicity and sensitivity to PI3Kalpha inhibitors. Science 366, 714–723 (2019). 10.1126/science.aaw9032

21 Schwartz, S. et al. Feedback suppression of PI3Kalpha signaling in PTEN-mutated tumors is relieved by selective inhibition of PI3Kbeta. Cancer Cell 27, 109–122 (2015). 10.1016/j.ccell.2014.11.008

22 Rodrik-Outmezguine, V. S. et al. mTOR kinase inhibition causes feedback-dependent biphasic regulation of AKT signaling. Cancer Discov 1, 248–259 (2011). 10.1158/2159-8290.CD-11-0085

23 Levina, A., Fleming, K. D., Burke, J. E. & Leonard, T. A. Activation of the essential kinase PDK1 by phosphoinositide-driven trans-autophosphorylation. Nat Commun 13, 1874 (2022). 10.1038/s41467-022-29368-4

24 Engelman, J. A. Targeting PI3K signalling in cancer: opportunities, challenges and limitations. Nat Rev Cancer 9, 550–562 (2009). 10.1038/nrc2664

25 He, Y. et al. Targeting PI3K/Akt signal transduction for cancer therapy. Signal Transduct Target Ther 6, 425 (2021). 10.1038/s41392-021-00828-5

26 Shaw, A. L. et al. ATP-competitive and allosteric inhibitors induce differential conformational changes at the autoinhibitory interface of Akt1. Structure 31, 343–354 e343 (2023). 10.1016/j.str.2023.01.007

27 Davies, B. R. et al. Preclinical pharmacology of AZD5363, an inhibitor of AKT: pharmacodynamics, antitumor activity, and correlation of monotherapy activity with genetic background. Mol Cancer Ther 11, 873–887 (2012). 10.1158/1535-7163.MCT-11-0824-T

28 Burnett, G. L. et al. Discovery of RMC-5552, a Selective Bi-Steric Inhibitor of mTORC1, for the Treatment of mTORC1-Activated Tumors. J Med Chem 66, 149–169 (2023). 10.1021/acs.jmedchem.2c01658

29 Li, J., Kim, S. G. & Blenis, J. Rapamycin: one drug, many effects. Cell Metab 19, 373–379 (2014). 10.1016/j.cmet.2014.01.001

30 Dowling, R. J. et al. mTORC1-mediated cell proliferation, but not cell growth, controlled by the 4E-BPs. Science 328, 1172–1176 (2010). 10.1126/science.1187532

31 Fingar, D. C. et al. mTOR controls cell cycle progression through its cell growth effectors S6K1 and 4E-BP1/eukaryotic translation initiation factor 4E. Mol Cell Biol 24, 200–216 (2004). 10.1128/MCB.24.1.200-216.2004

32 Aviner, R. The science of puromycin: From studies of ribosome function to applications in biotechnology. Comput Struct Biotechnol J 18, 1074–1083 (2020). 10.1016/j.csbj.2020.04.014

33 O’Reilly, K. E. et al. mTOR inhibition induces upstream receptor tyrosine kinase signaling and activates Akt. Cancer Res 66, 1500–1508 (2006). 10.1158/0008-5472.CAN-05-2925

34 Holderfield, M. et al. Concurrent inhibition of oncogenic and wild-type RAS-GTP for cancer therapy. Nature 629, 919–926 (2024). 10.1038/s41586-024-07205-6

35 Kitai, H. et al. Combined inhibition of KRAS(G12C) and mTORC1 kinase is synergistic in non-small cell lung cancer. Nat Commun 15, 6076 (2024). 10.1038/s41467-024-50063-z

36 Schulze, C. J. et al. Chemical remodeling of a cellular chaperone to target the active state of mutant KRAS. Science 381, 794–799 (2023). 10.1126/science.adg9652

37 Chandarlapaty, S. et al. AKT inhibition relieves feedback suppression of receptor tyrosine kinase expression and activity. Cancer Cell 19, 58–71 (2011). 10.1016/j.ccr.2010.10.031

38 Lee, B. J. et al. Selective inhibitors of mTORC1 activate 4EBP1 and suppress tumor growth. Nat Chem Biol 17, 1065–1074 (2021). 10.1038/s41589-021-00813-7

39 Schram, A. M. et al. The Bi-steric, mTORC1-Selective Inhibitor, RMC-5552, in Advanced Solid Tumors: A Phase 1 Trial. Clin Cancer Res (2025). 10.1158/1078-0432.CCR-25-2112

40 Mattar, M. et al. Establishing and Maintaining an Extensive Library of Patient-Derived Xenograft Models. Front Oncol 8, 19 (2018). 10.3389/fonc.2018.00019

41 Yi, S. A., Sepic, S., Schulman, B. A., Ordureau, A. & An, H. mTORC1-CTLH E3 ligase regulates the degradation of HMG-CoA synthase 1 through the Pro/N-degron pathway. Mol Cell 84, 2166–2184 e2169 (2024). 10.1016/j.molcel.2024.04.026

