## Supplementary information for "Coexistent *PTEN* and *PIK3CA* alterations hyperactivate mTORC1 signaling in endometrial cancers and cause their selective sensitivity to mTORC1 inhibition"

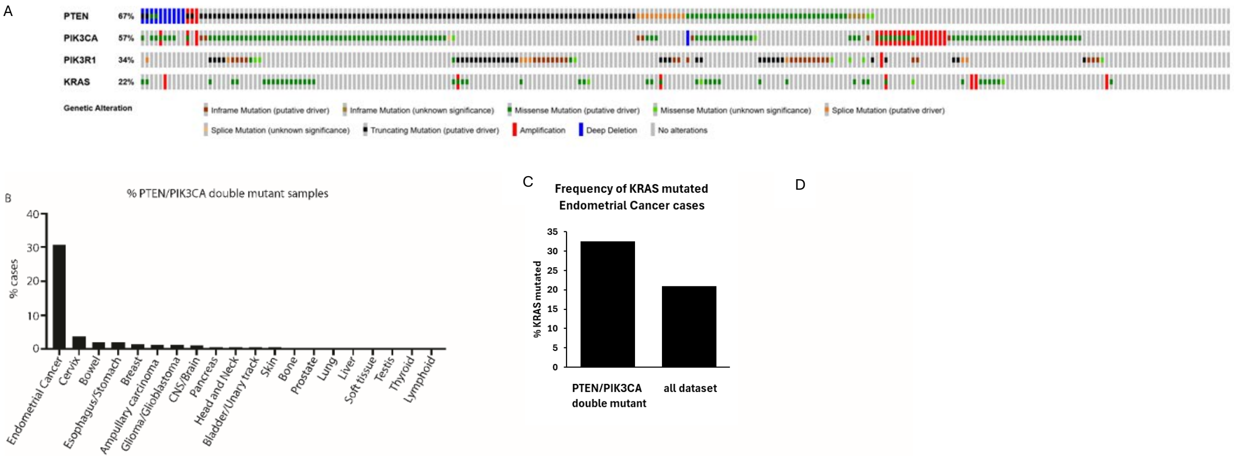


**Supplementary Figure 1: Analysis of patient-derived database using the cBioPortal and FMI[7-9].** (A) Analysis of EC patients-derived Uterine Corpus Endometrial Carcinoma samples (TCGA, Firehose Legacy dataset, cBioPortal for Cancer Genomics) showing the prevalence of *PTEN*, *PIK3CA*, *PIK3R1* and *KRAS* alterations. (B) cBioPortal database analysis of *PIK3CA* and *PTEN* mutations, showing the percentages of *PTEN* and *PIK3CA* coexistent mutations in various cancer types. (C) Frequency of *KRAS* mutations in endometrial cancer patient samples with *PTEN* and *PIK3CA* co-alterations compared with all endometrial cancer dataset (derived from TCGA, Firehose Legacy dataset, cBioPortal for Cancer Genomics) (D) Frequency of *KRAS* short variant mutations in endometrial cancer patient samples with or without *PTEN* and *PIK3CA* co-alterations (derived from Foundation Medicine Insights (version MI20240507))

| Cell line | *PTEN* | *PIK3CA* |
|---|---|---|
| **KLE** | WT | WT |
| **MFE280** | WT | H1047Y; I391M |
| **AN3CA** | R130Qfs | WT |
| **MFE296** | R130Q, N323fs | P539R, I20M |
| **HEC6** | V290fs, V85fs | R108H, C420fs |
| **High-grade uterine carcinoma, PDX** | M134del, R173H | E542K |
| **Endometrioid adenocarcinoma, PDX** | L98Qfs | E545K |
| **Endometrioid adenocarcinoma, PDX** | G290Sfs | E542A, R38C |
| **NOU-1** | No mRNA expression | R38H |

**Supplementary Table 1:** *PTEN* and *PIK3CA* status in various EC models [17]

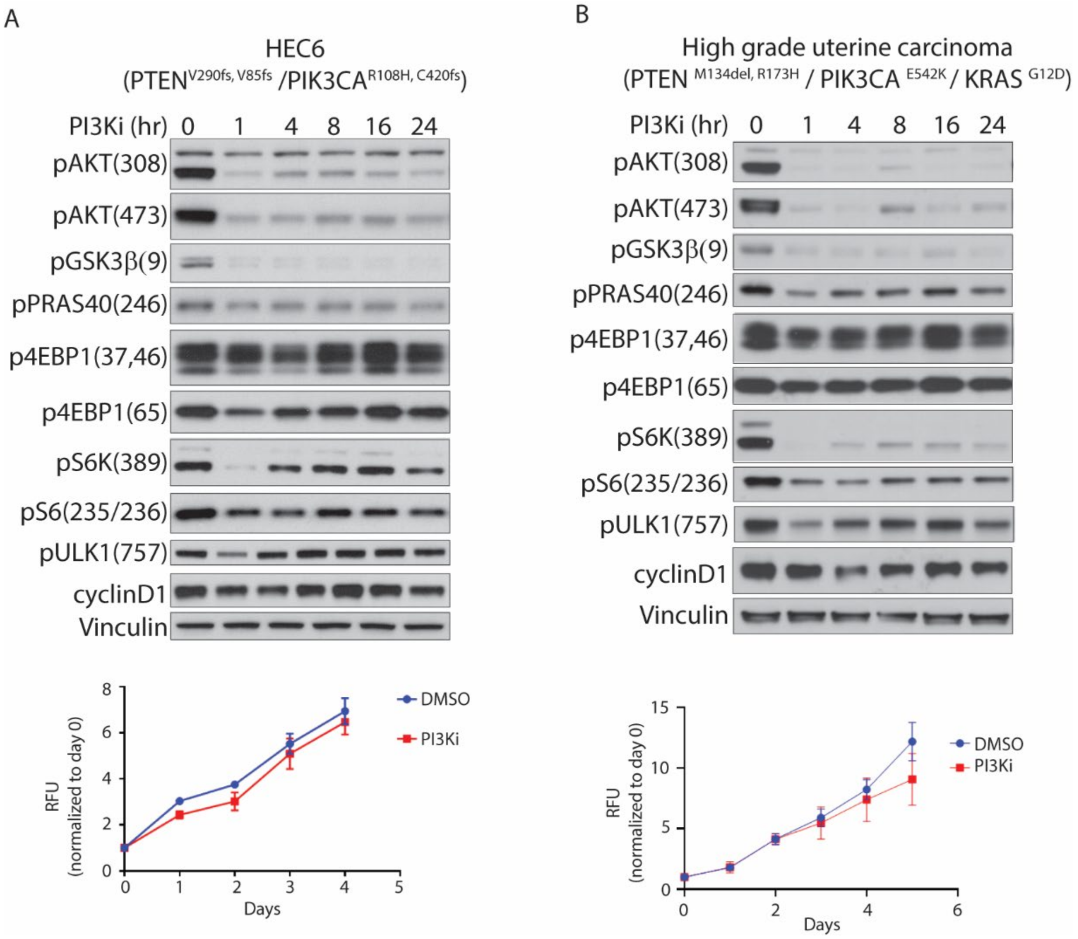


**Supplementary Figure 2: EC tumor cells with coexistent *PTEN* inactivation and activated *PIK3CA* mutants are insensitive to PI3K inhibition.** HEC6 ($PTEN^{V290fs,\ V85fs}$/$PIK3CA^{R108H,\ C420fs}$) cells (A) or high-grade uterine carcinoma ($PTEN^{M134del,\ R173H}$ / $PIK3CA^{E542K}$ / $KRAS^{G12D}$) PDX-derived cells (B) were treated with combination of BYL719 (1μM) and AZD8186 (250nM) for the indicated time points. Upper panels: Western Blots depict PI3K/mTOR signaling output. Lower panels: cell viability measured by alamarBlue assay.

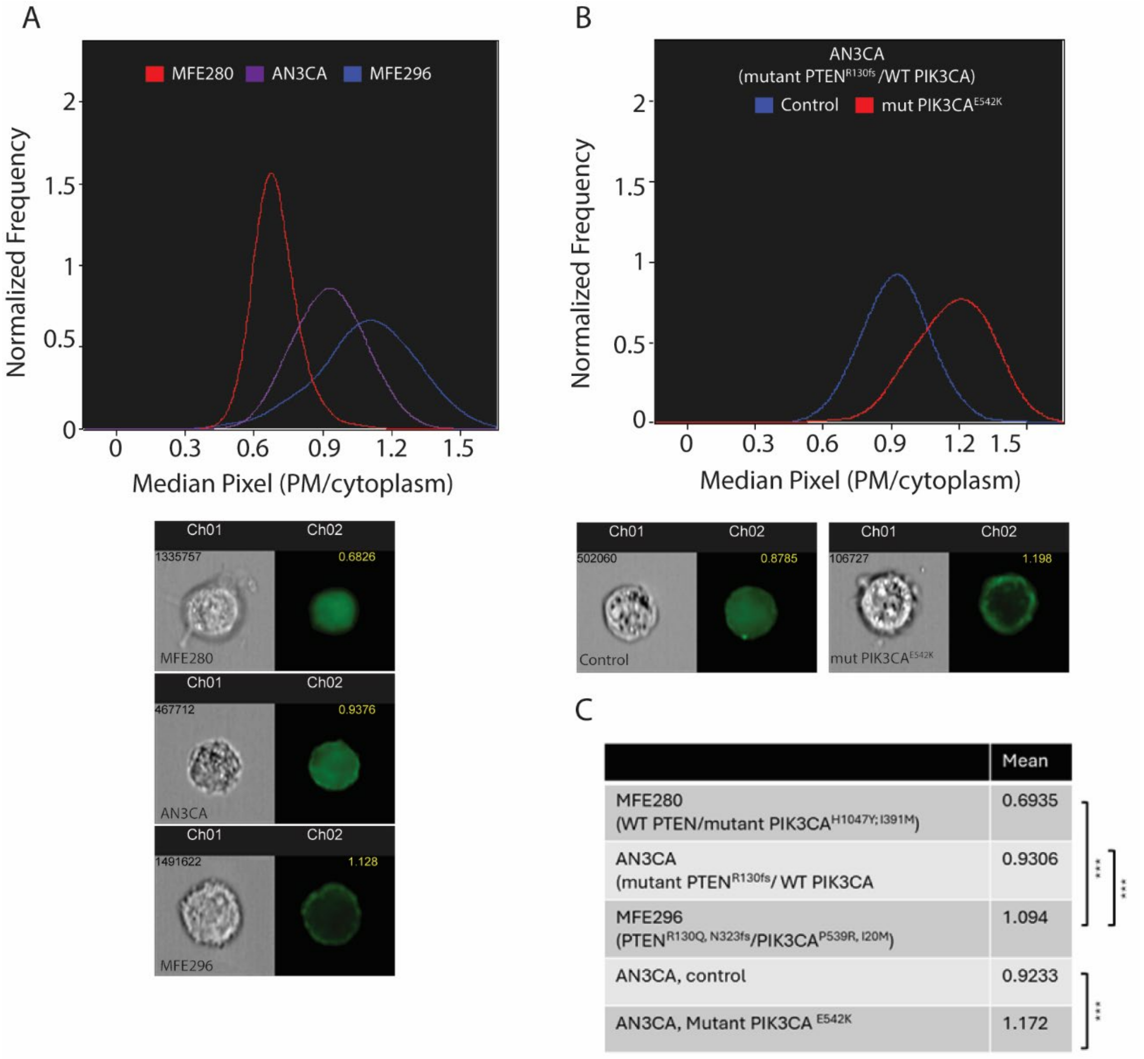


**Supplementary Figure 3: PIP3 levels are higher in the *PTEN/PIK3CA* double mutant EC cells**. The pcDNA3-AKT-PH-GFP vector [22] (GFP-PH$^{AKT}$) was transfected into AN3CA, MFE280 and MFE296 cells (A) or transfected to AN3CA together with mutant *PIK3CA*$^{E542K}$ over-expression plasmid (B). After 48 hours cells were analyzed for the GFP-PH$^{AKT}$ probe translocation to the plasma membrane (PM) by ImageStreamX, using masking tool. The histograms illustrate the distribution of cells based on the ratio scores of the mean GFP pixel intensity in the plasma membrane relative to the cytoplasm. The images in lower panels represent sample cells displaying the most common ratio within the indicated populations (peak of the histograms). MFE280 n= 92, AN3CA n= 1432, MFE296 n= 1367, AN3CA control n= 557, AN3CA mutant *PIK3CA*$^{E542K}$ n= 668. Experiments with AN3CA isogenic cells and MFE296 were repeated three times, and those with MFE280 and AN3CA parental cells was repeated twice. Similar results were obtained, and representative results are shown. Statistical significance was calculated by pairwise Welch t-test ***p-value <0.001

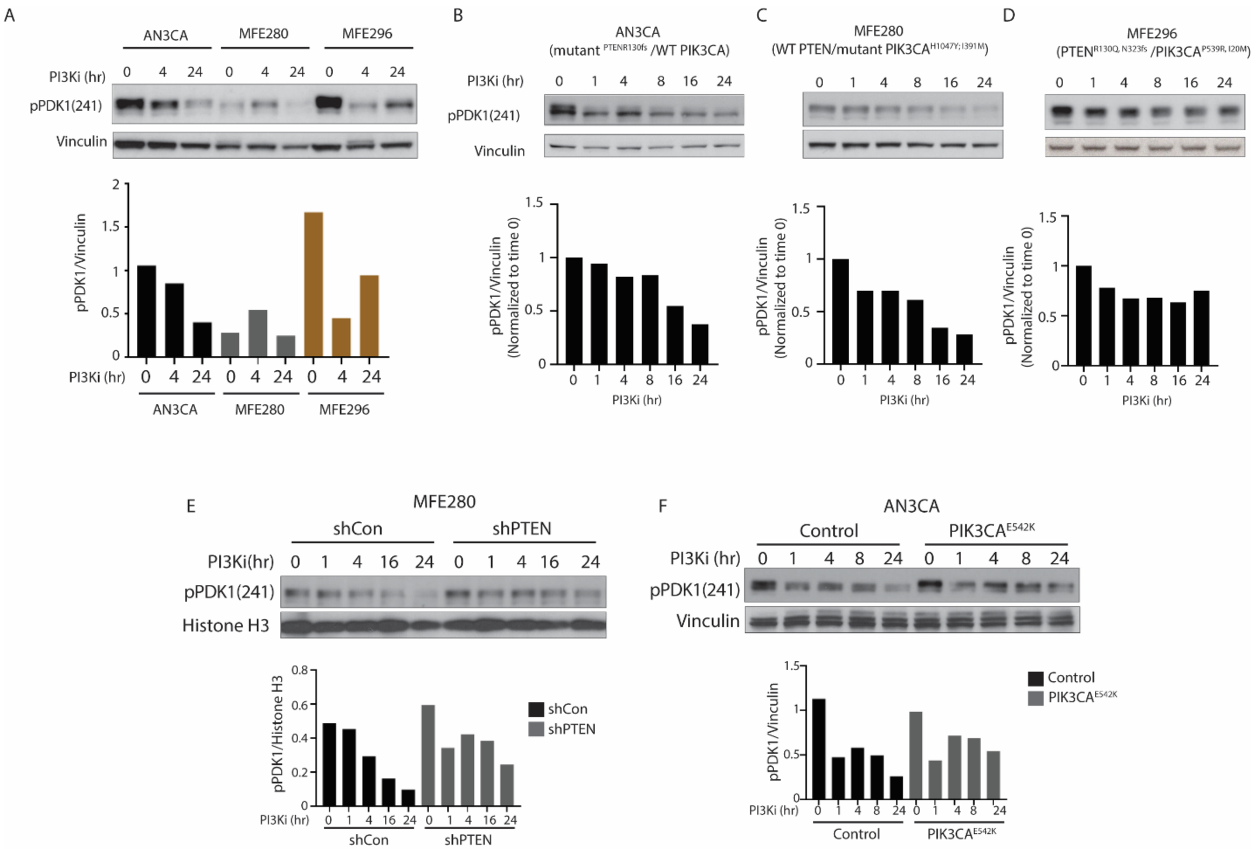


**Supplementary Figure 4: PDK1 phosphorylation at S241 is higher in the *PTEN/PIK3CA* coexistent cells (MFE296) compared with single mutant cells (AN3CA or MFE280).** Western Blots presenting the phosphorylated PDK1 at S241 in AN3CA, MFE280 and MFE296 (A-D) and in MFE280 (E) and AN3CA (F) isogenic cells, upon PI3K inhibition by BYL719(1μM) +AZD8186(250nM) treatment for the indicated time points. The experiments in panels B–F are the same as those described in Figure 1.

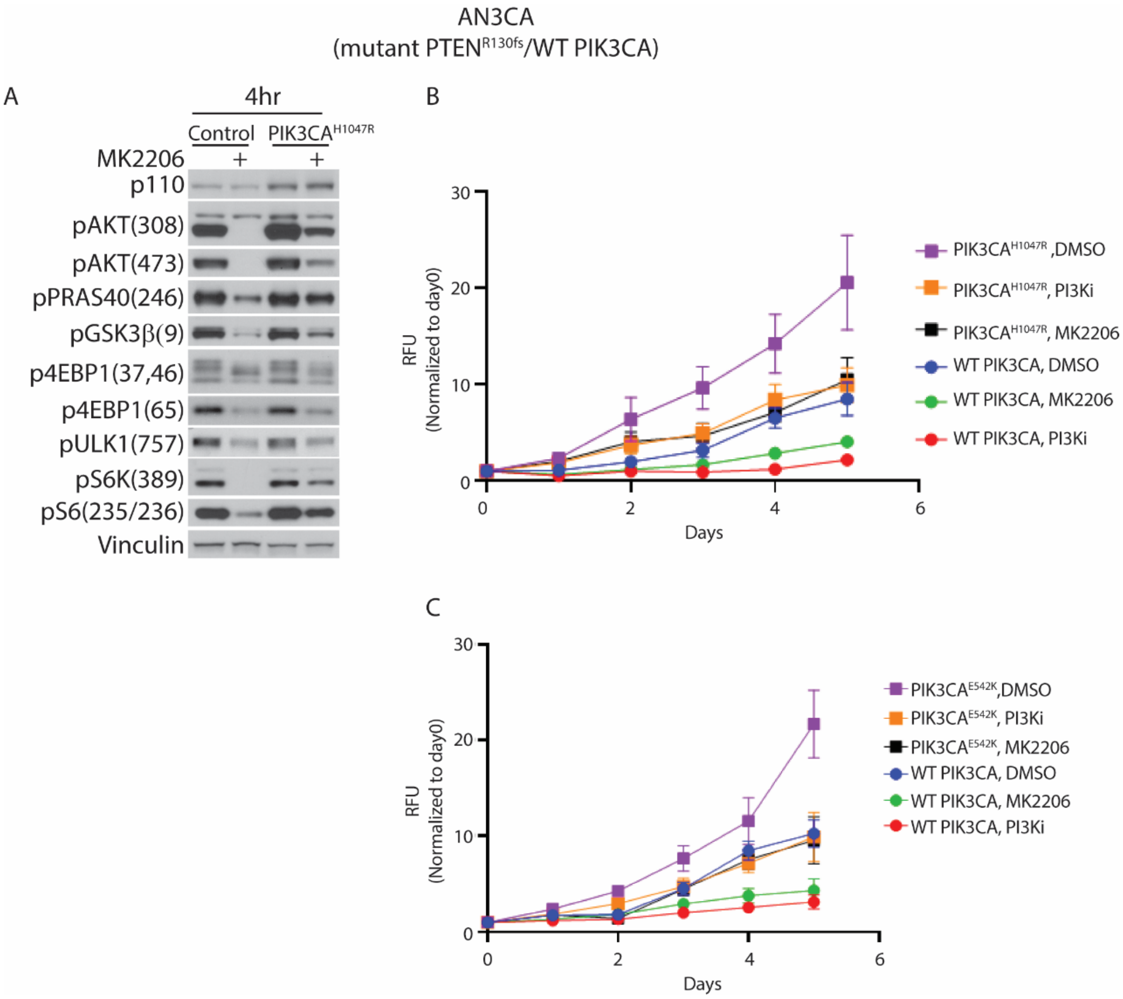


**Supplementary Figure 5: *PTEN/PIK3CA* double mutant isogenic AN3CA cells are insensitive to AKT inhibition**. AN3CA (mutant *PTEN*$^{R130fs}$ /WT *PIK3CA*) cells were transiently transfected with mutant *PIK3CA*$^{H1047R}$. (A) After 24 hours, cells were treated with 1 µM MK2206 for 4 hours. Immunoblots depicting PI3K/mTOR signaling output. (B) After 24 hours, cells were treated with either 1 µM MK2206 or a combination of 1µM BYL-719 + 250nM AZD8186 (PI3Ki) for the indicated time points. Cell proliferation was assessed by alamarBlue assay (C) AN3CA (mutant *PTEN*$^{R130fs}$ /WT *PIK3CA*) cells were transiently transfected with mutant *PIK3CA*$^{E542K}$. After 24 hours, cells were treated with either 1 µM MK2206 or a combination of 1µM BYL-719 + 250nM AZD8186 (PI3Ki) for the indicated time points. Cell proliferation was assessed by alamarBlue assay. Growth curves for MK2206 and DMSO are those shown in Figure 2F.

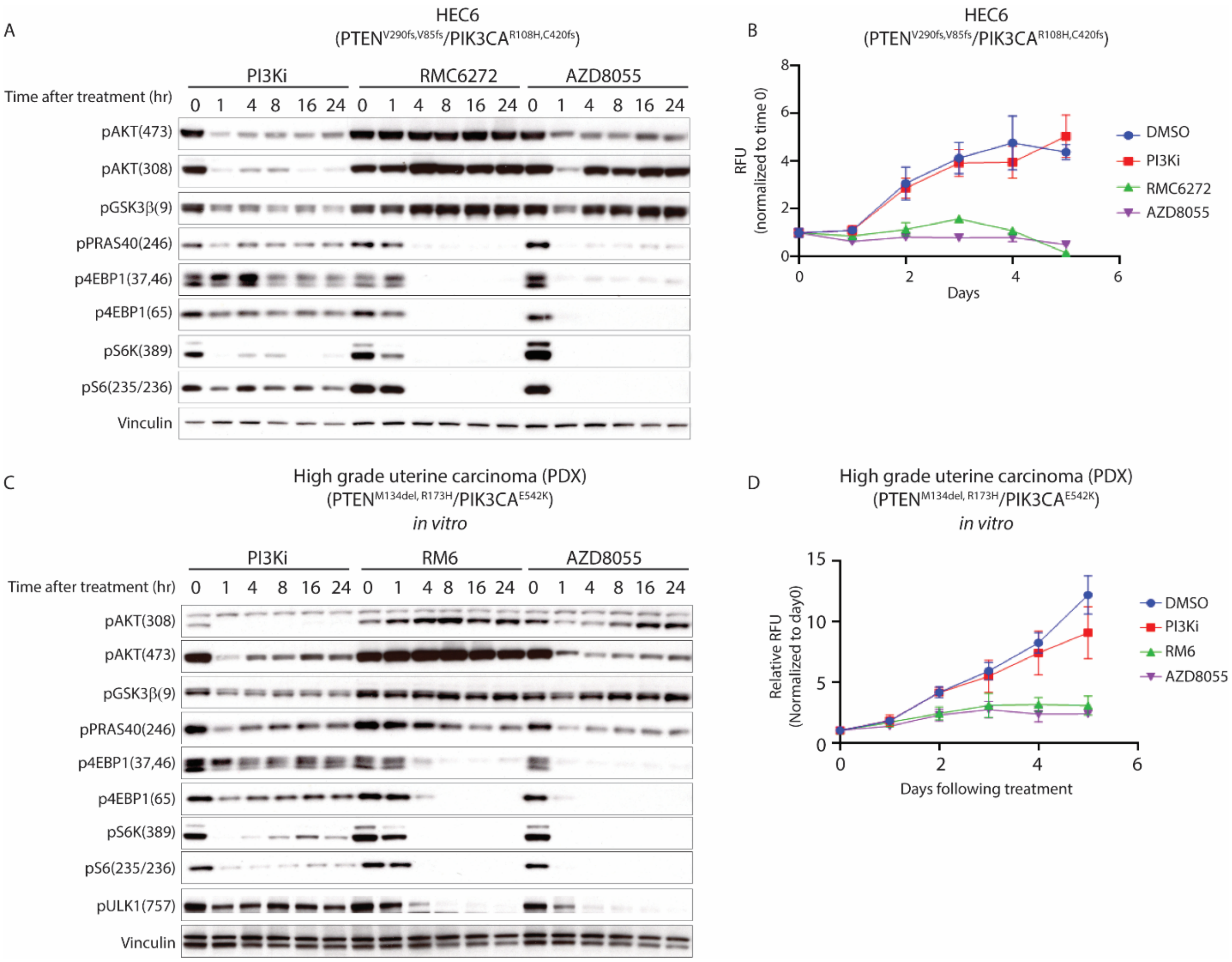


**Supplementary Figure 6: *PTEN/PIK3CA* double mutant EC cells are dependent on mTORC1**. HEC6 (*PTEN*$^{V290fs,\ V85fs}$ /*PIK3CA*$^{R108H,\ C420fs}$) (A-B) and the High-grade Uterine carcinoma PDX-derived cells (*PTEN*$^{M134del,\ R173H}$ / *PIK3CA*$^{E542K}$ /*KRAS*$^{G12D}$) (C-D) were treated *in vitro* with either of the following inhibitors: BYL-719 (1µM) + AZD8186 (250nM) (PI3Ki), RMC-6272 (500pM) or AZD8055 (500nM) for the indicated time points. Cell growth was assessed by alamarBlue assay (B and D), and signaling output was measured by western blot using specific antibodies (A and C).

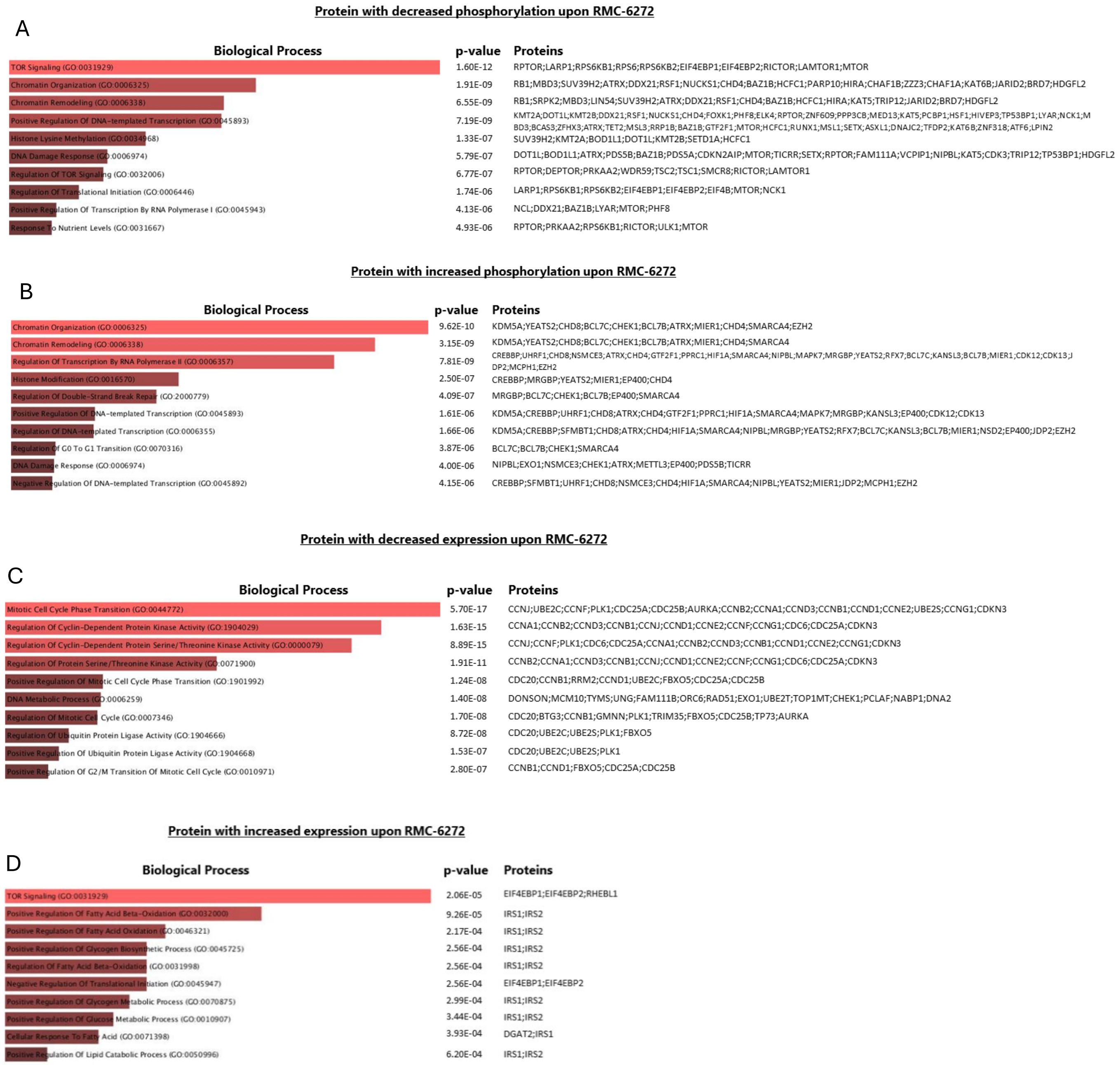


**Supplementary Figure 7: Biological processes enriched in protein sets that were changed upon RMC-6272 treatment.** MFE296 cells (*PTEN*$^{R130Q,\ N323fs}$ /*PIK3CA*$^{P539R,\ I20M}$) were treated with either a combination of PI3K inhibitors BYL719 (1µM) and AZD8186 (250nM), RMC-6272 (500pM) or Torin1 (250nM) for 16 hours followed by global unbiased proteomic analysis by mass-spectrometry. Proteins whose expression or phosphorylation was significantly changed upon treatment were defined as a fold change of ≥2 or ≤0.5 with a p-value ≤0.01. Enrichment analysis was performed using the suite of gene set enrichment analysis tools, Enrichr [42-44]. The figure shows the 10 most enriched annotations found in the 4 defined protein groups, their p-value and protein lists included in the groups. (A) proteins with decreased phosphorylation (B) proteins with increased phosphorylation (C) proteins with decreased total expression (D) proteins with increased total expression.

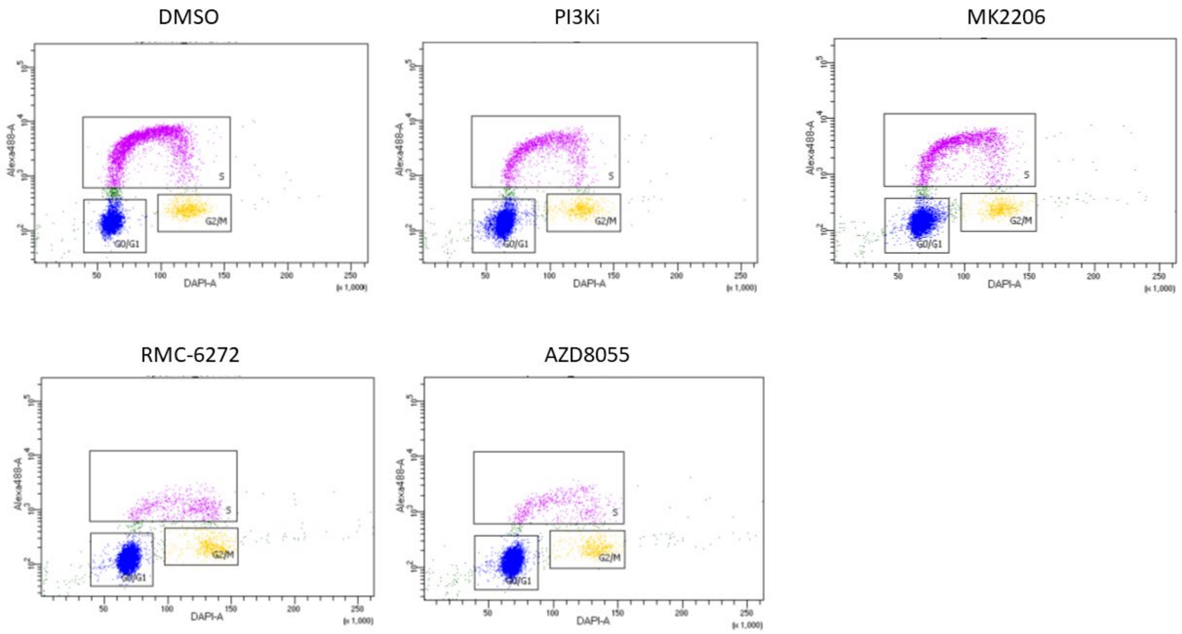


**Supplementary Figure 8: FACS dot plots corresponding to Figure 4F.** MFE296 cells were treated with either PI3K inhibitors BYL719 (1μM) and AZD8186 (250nM), MK2206(1μM), AZD8055 (500nM) or with RMC-6272 (500pM) for 24 hours. Cell cycle states were detected using EdU-DAPI-based flow cytometry. Dot plots of representative experiment are shown.

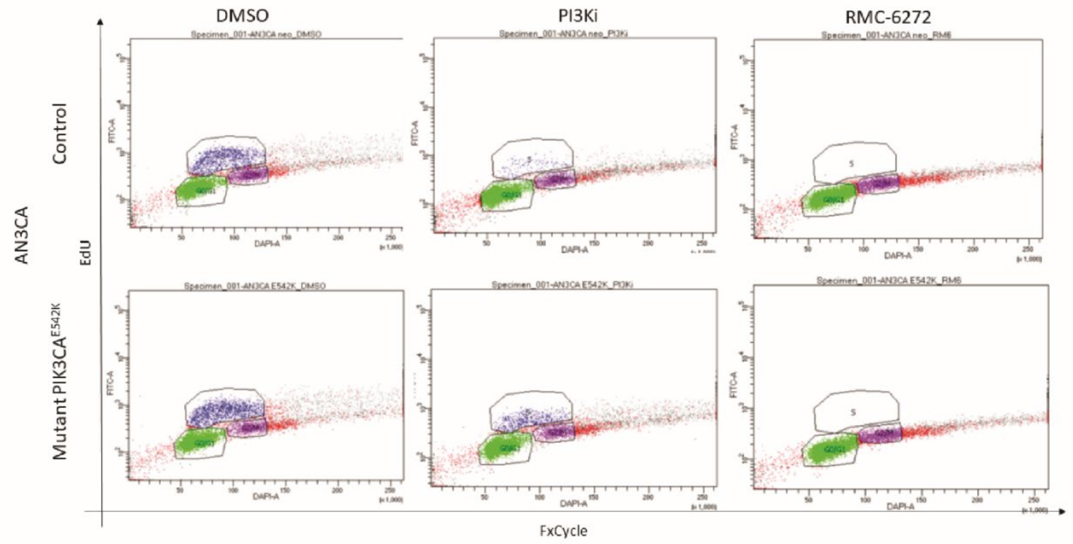


**Supplementary Figure 9: FACS dot plots corresponding to Figure 4G** AN3CA cells were transiently transfected with mutant *PIK3CA*$^{E452K}$, and after 24 hours were treated with either PI3K inhibitors BYL719 (1μM) and AZD8186 (250nM), or with RMC-6272 (500pM) for additional 24 hours. Cell cycle states were detected using EdU-DAPI-based flow cytometry. Dot plots of representative experiment are shown.

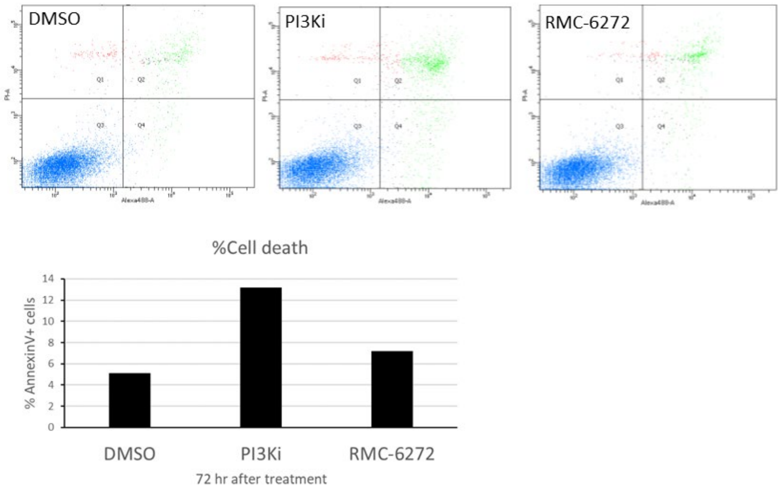


**Supplementary Figure 10: Apoptosis was not significantly induced by mTOR inhibition.** MFE296 cells were treated with either PI3K inhibitors: BYL719(1μM)+ AZD8186(250nM), or RMC-6272 (500pM), for 72 hours, followed by Annexin V and PI staining. Cell percentages were analyzed by flow cytometry.

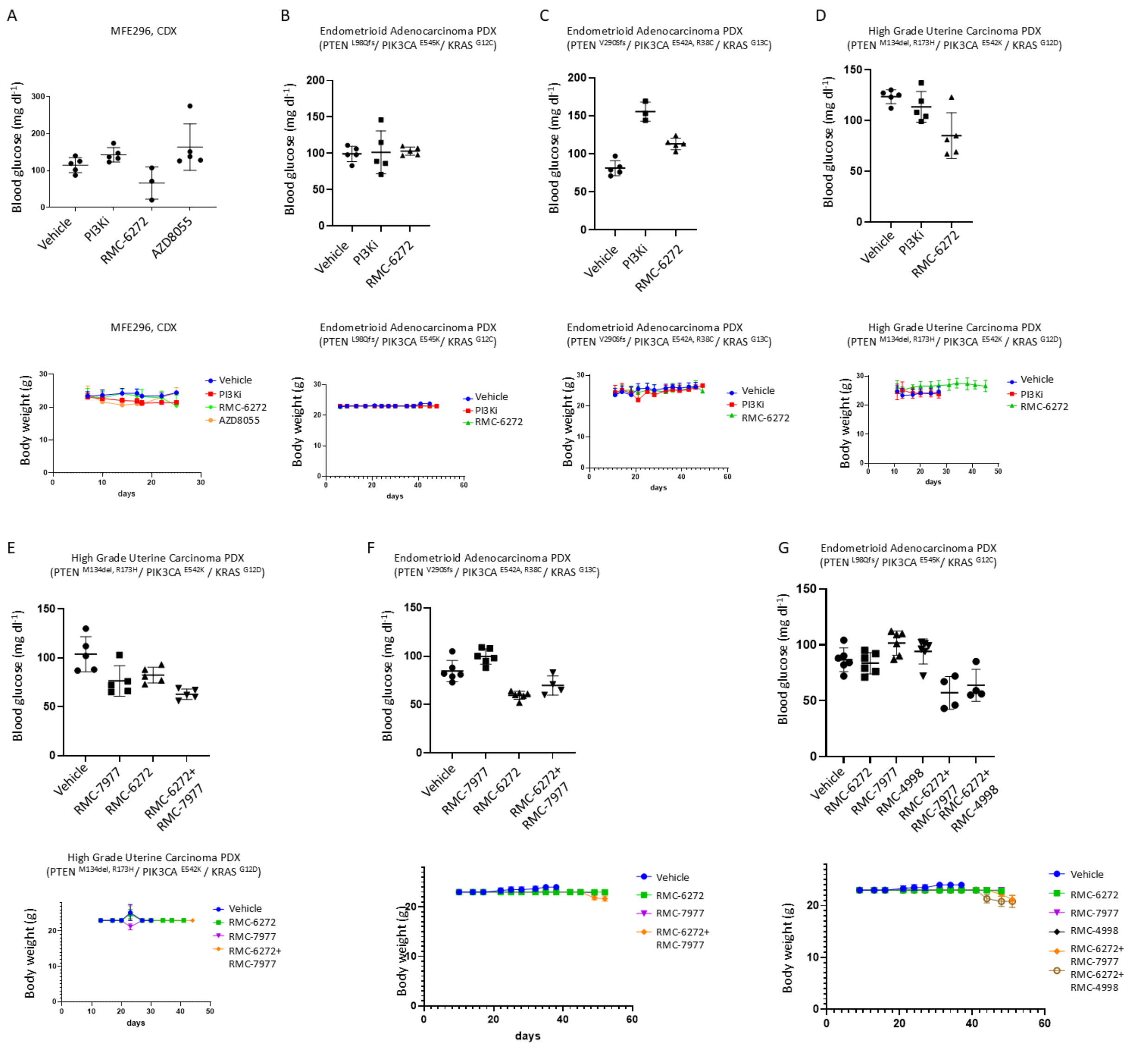

**Supplementary Figure 11:** Blood glucose levels (upper panels) and body weight (lower panels) detected in the mice of *in vivo* experiments presented in Figure 5 (A-D) and 6 (E-G).

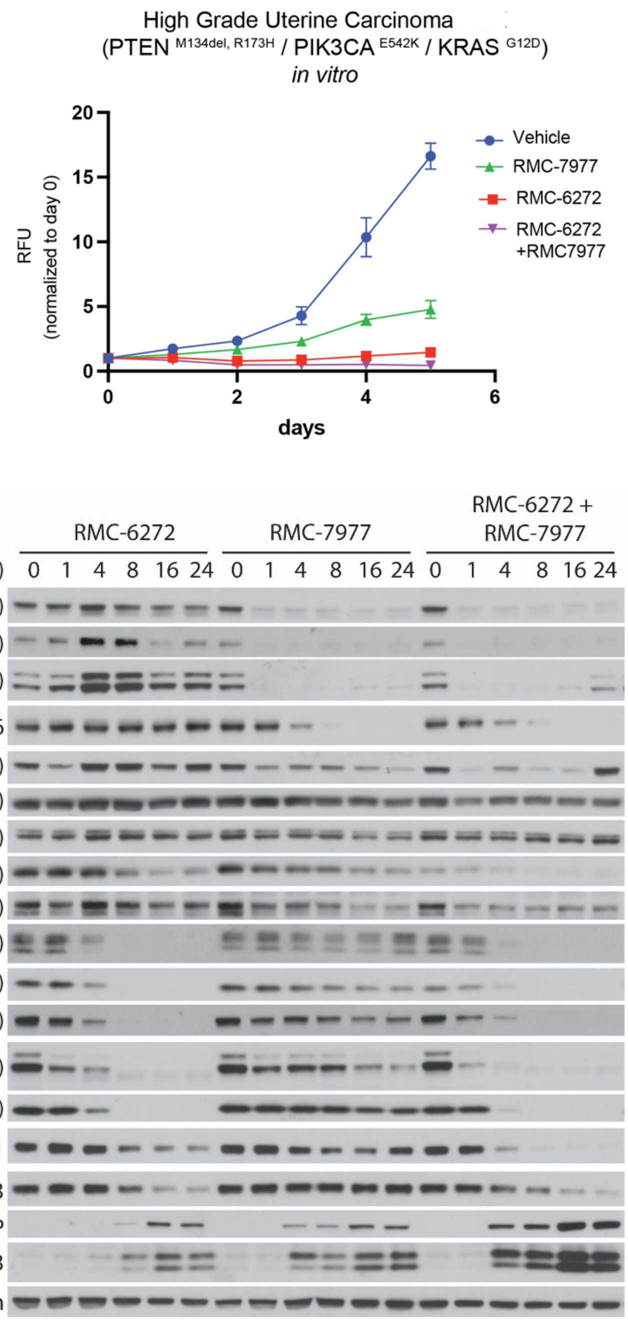


**Supplementary Figure 12:** High Grade Uterine Carcinoma PDX derived cells (*PTEN*$^{M13del,R173}$ / *PIK3C* $^{E542K}$ / *KRAS*$^{G12D}$) were cultured *in vitro* and treated with RMC-6272 (500pM), RMC-7977 (100nM) or combination of both inhibitors for the indicated time points. Upper panel: Cell growth was assessed by Presto Blue assay. Lower panel: PI3K/mTOR and ERK and signaling output was measured by western blot using specific antibodies.

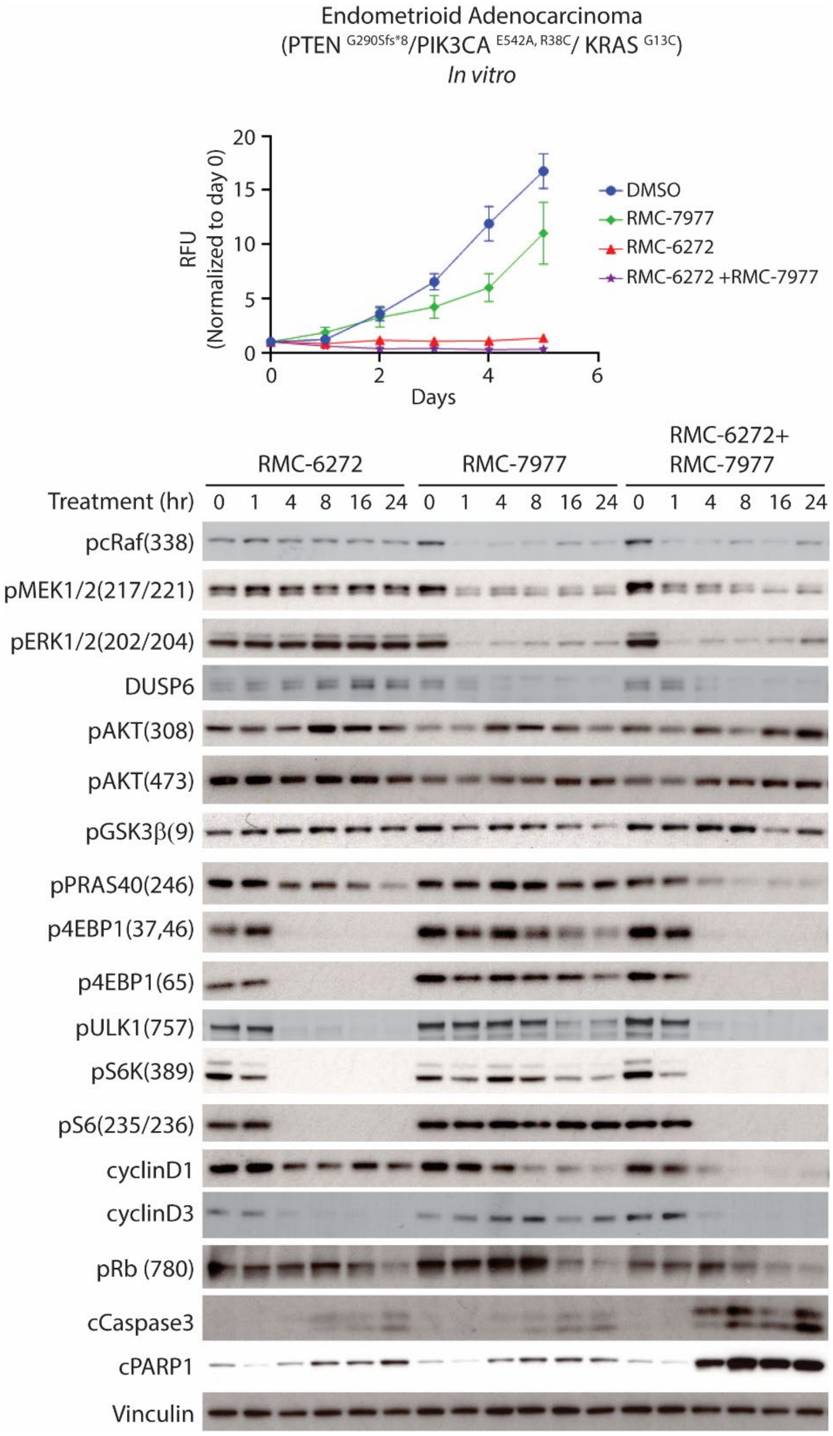


**Supplementary Figure 13:** Endometrioid adenocarcinoma PDX derived cells (*PTEN*$^{V290Sfs}$ / *PIK3CA*$^{E542A, R38C}$ / *KRAS*$^{G13C}$) were cultured *in vitro* and treated with RMC-6272 (500pM), RMC-7977 (100nM) or combination of both inhibitors for the indicated time points. Upper panel: Cell growth was assessed by Presto Blue assay. Lower panel: PI3K/mTOR and ERK and signaling output was measured by western blot using specific antibodies.

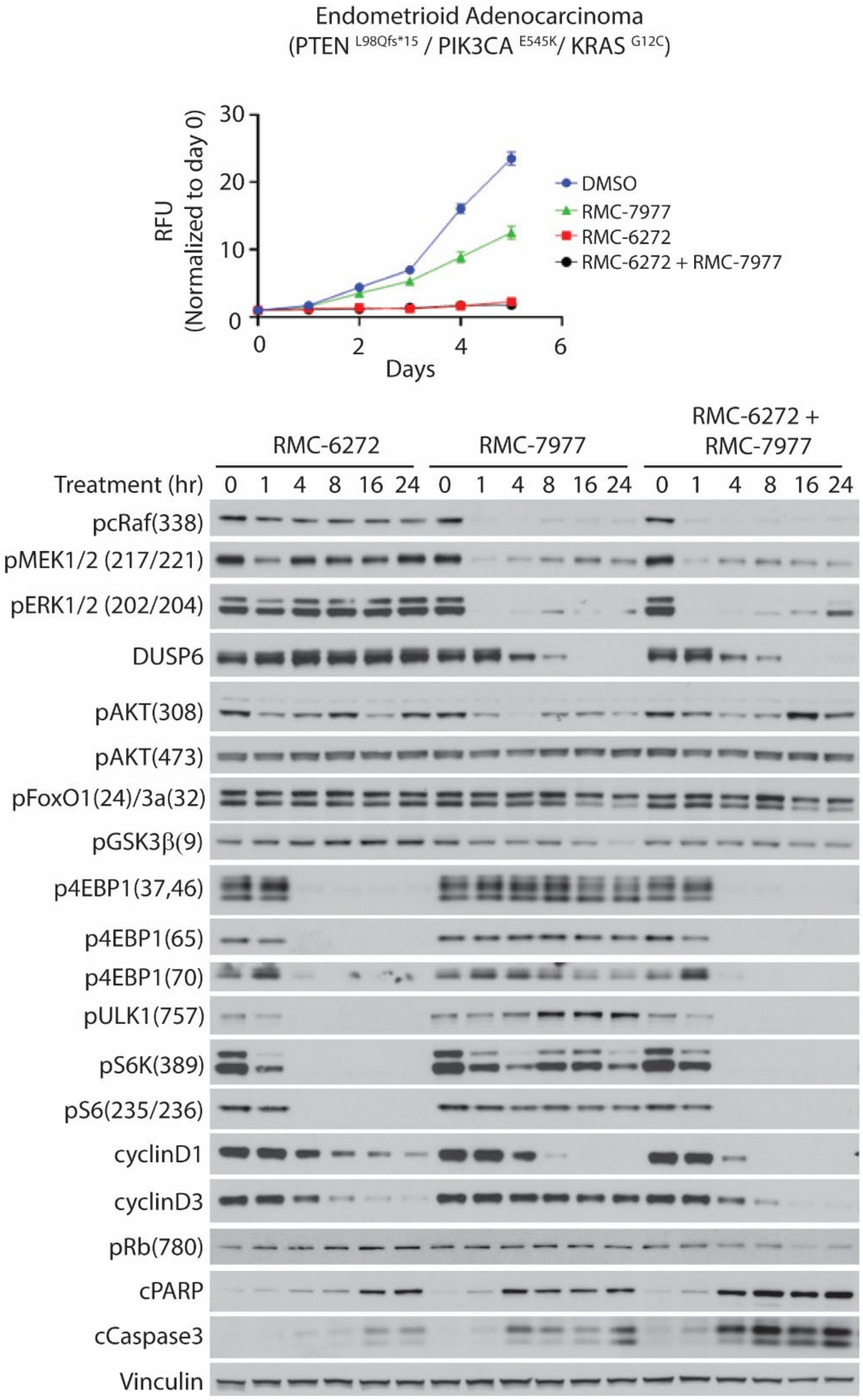


**Supplementary Figure 14:** Endometrioid adenocarcinoma PDX derived cells (*PTEN* $^{L98Qfs}$/ *PIK3CA* $^{E545K}$/ *KRAS* $^{G12C}$) were cultured *in vitro* and treated with RMC-6272 (500pM), RMC-7977 (100nM) or combination of both inhibitors for the indicated time points. Upper panel: Cell growth was assessed by Presto Blue assay. Lower panel: PI3K/mTOR and ERK and signaling output was measured by western blot using specific antibodies.

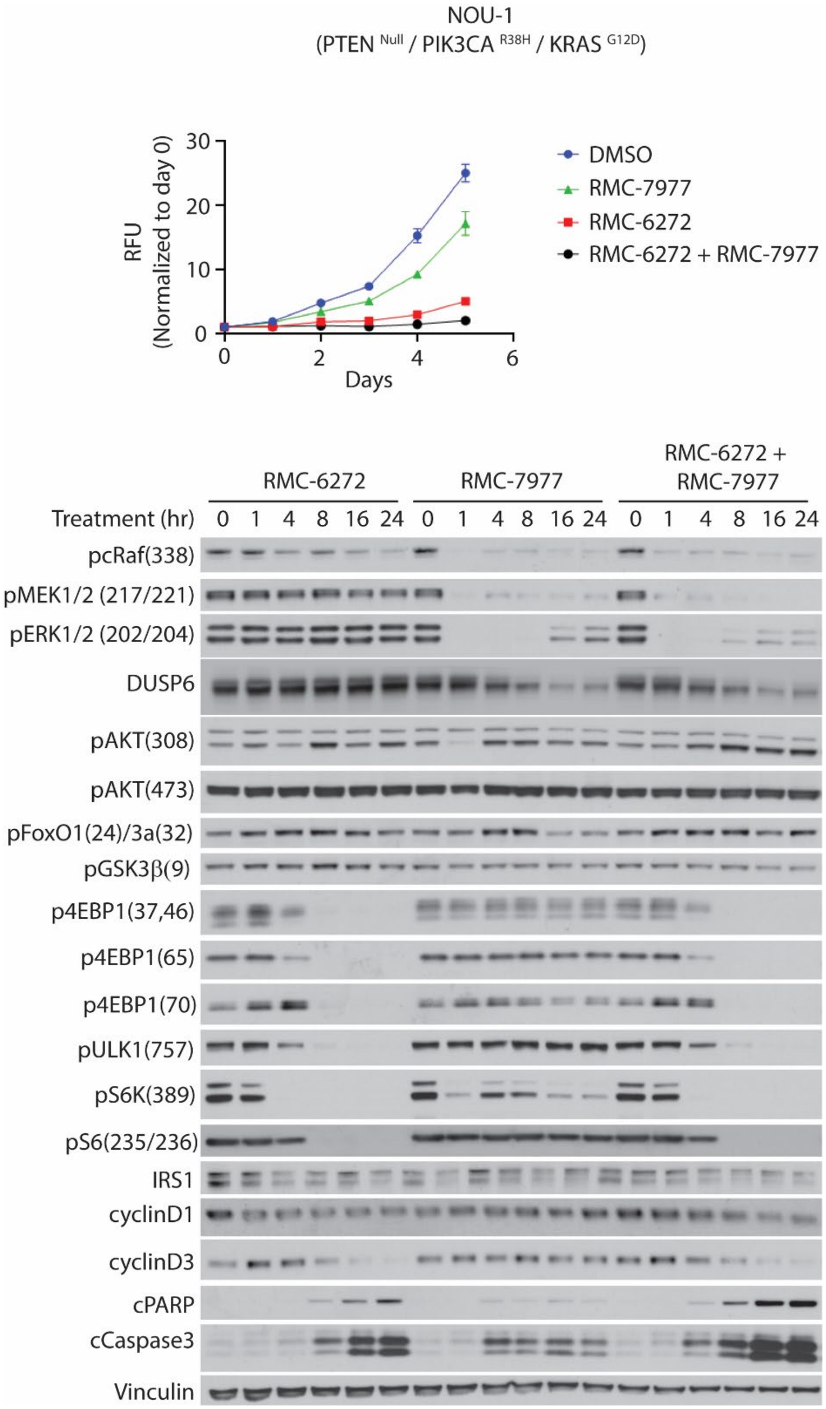


**Supplementary Figure 15:** NOU1 cell line (*PTEN* $^{Null}$*/ PIK3CA* $^{R38H}$*/ KRAS* $^{G12D}$) was cultured *in vitro* and treated with RMC-6272 (500pM), RMC-7977 (100nM) or combination of both inhibitors for the indicated time points. Upper panel: Cell growth was assessed by Presto Blue assay. Lower panel: PI3K/mTOR and ERK and signaling output was measured by western blot using specific antibodies.

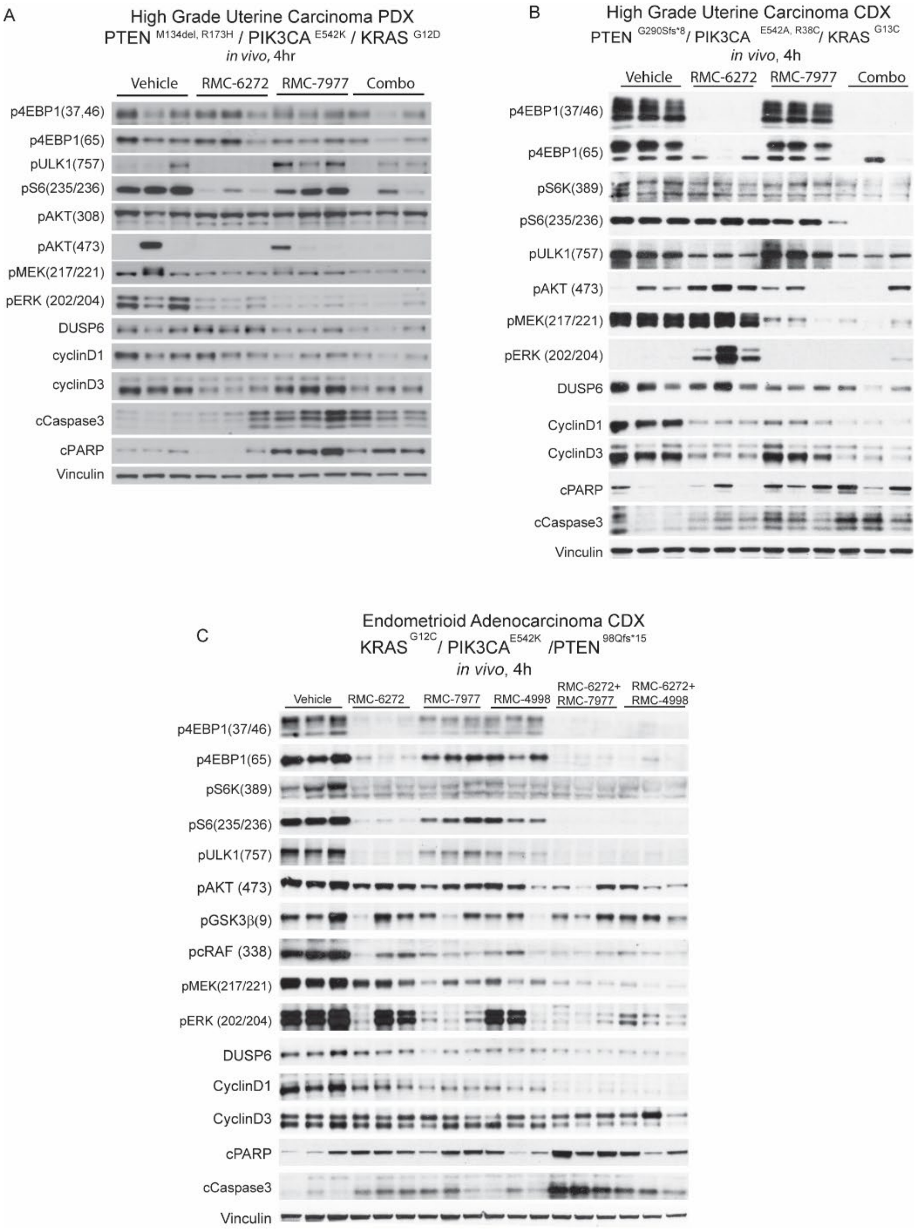


**Supplementary Figure 16:** (A) Mice bearing high grade uterine carcinoma (*PTEN* $^{M134del,\ R173H}$ / *PIK3CA* $^{E542K}$ / *KRAS* $^{G12D}$) patient-derived xenograft (PDX) tumors were treated *in vivo* with RMC-6272 (3 mg/kg i.p. QW), RMC-7977 (25mg/kg p.o. 3 day/week) or their combination. Immunoblots depict RAS/PI3K/mTOR signaling output after 4 hours of treatment. (B) Endometrioid adenocarcinoma (*PTEN* $^{V290Sfs}$ / *PIK3CA* $^{E542A,\ R38C}$ / *KRAS* $^{G13C}$) PDX cells were cultured *in vitro* and subsequently implanted into mice to establish cell-derived xenograft (CDX) tumors. Mice were treated *in vivo* with RMC-6272 (3 mg/kg i.p. QW), RMC-7977 (25mg/kg p.o. 3 day/week) or their combination. Immunoblots depict RAS/PI3K/mTOR signaling output after 4 hours of treatment. (C) Endometrioid adenocarcinoma (*PTEN* $^{L98Qfs}$/ *PIK3CA* $^{E545K}$/ *KRAS* $^{G12C}$) PDX cells were cultured *in vitro* and subsequently implanted into mice to establish cell-derived xenograft (CDX) tumors. Mice were treated *in vivo* with RMC-6272 (3 mg/kg i.p. QW), RMC-7977 (25mg/kg p.o. 3 day/week), RMC-4998 (80mg/kg p.o. QDx5) or either RMC-6272+ RMC-7977 or RMC-6272+ RMC-4998 combinations. Immunoblots depict RAS/PI3K/mTOR signaling output after 4 hours of treatment.

### References


1 Cerami, E. *et al.* The cBio cancer genomics portal: an open platform for exploring multidimensional cancer genomics data. *Cancer Discov* **2**, 401-404 (2012). https://doi.org:10.1158/2159-8290.CD-12-0095

2 de Bruijn, I. *et al.* Analysis and Visualization of Longitudinal Genomic and Clinical Data from the AACR Project GENIE Biopharma Collaborative in cBioPortal. *Cancer Res* **83**, 3861-3867 (2023). https://doi.org:10.1158/0008-5472.CAN-23-0816

3 Gao, J. *et al.* Integrative analysis of complex cancer genomics and clinical profiles using the cBioPortal. *Sci Signal* **6**, pl1 (2013). https://doi.org:10.1126/scisignal.2004088

4 Weigelt, B., Warne, P. H., Lambros, M. B., Reis-Filho, J. S. & Downward, J. PI3K pathway dependencies in endometrioid endometrial cancer cell lines. *Clin Cancer Res* **19**, 3533-3544 (2013). https://doi.org:10.1158/1078-0432.CCR-12-3815

5 Rodrik-Outmezguine, V. S. *et al.* mTOR kinase inhibition causes feedback-dependent biphasic regulation of AKT signaling. *Cancer Discov* **1**, 248-259 (2011). https://doi.org:10.1158/2159-8290.CD-11-0085

6 Chen, E. Y. *et al.* Enrichr: interactive and collaborative HTML5 gene list enrichment analysis tool. *BMC Bioinformatics* **14**, 128 (2013). https://doi.org:10.1186/1471-2105-14-128

7 Kuleshov, M. V. *et al.* Enrichr: a comprehensive gene set enrichment analysis web server 2016 update. *Nucleic Acids Res* **44**, W90-97 (2016). https://doi.org:10.1093/nar/gkw377

8 Xie, Z. *et al.* Gene Set Knowledge Discovery with Enrichr. *Curr Protoc* **1**, e90 (2021). https://doi.org:10.1002/cpz1.90

9 Foundation Medicine insights (version MI20240507) [cited 2025 Aug 15]. Available from: https://foundationmedicine.com/service/genomic-data-solutions.
